# Understanding the mechanism of *Trikatu* in type 2 diabetes mellitus and lipid-related metabolic disorders: A network pharmacology approach

**DOI:** 10.1101/2022.06.22.496819

**Authors:** Varun Chhabra, Mohit Wadhawan, Amit Katiyar, Bharat Krushna Khuntia, Vandna Sharma, Shubhangi Rathore, Punit kaur, Gautam Sharma

## Abstract

**Objective:** Trikatu is an Indian polyherbal formulation comprising three herbs, i.e., Zingiber officinale, Piper longum, and Piper nigrum. It is traditionally used to treat metabolic disorders such as type 2 diabetes mellitus (T2DM), dyslipidemia, and obesity. However, its mechanism of action remains unknown. This study aims to explore the underlying therapeutic mechanism of Trikatu in T2DM and lipid metabolic disorders using network pharmacology (NP).

**Methods:** Trikatu phytochemicals were retrieved from various databases and screened on the basis of druglikeness and oral bioavailability (>30%) score. Putative targets of the bioactive phytochemicals were identified using TargetNet, Similarity Ensemble Approach, and Swiss Target Prediction databases. Protein-protein interaction (PPI) network of overlapping targets of phytochemicals and metabolic disorders was constructed using NetworkAnalyst 3.0. The Bioactive Phytochemical-Target-Pathway (BP-T-P) network was constructed using cytoscape v3.8.2, and the key targets of Trikatu were analyzed by Gene Ontology (GO), and Kyoto Encyclopedia of Genes and Genomes (KEGG) enrichment.

**Results:** Twenty bioactive phytochemicals and 102 targets of Trikatu were identified. PPI network and enrichment analysis showed that 51 targets of Trikatu phytochemicals such as RXRA, STAT3 and ESR1, GSK3B, TNF, NOS2/3 regulate pathways like insulin resistance, steroid hormone biosynthesis, regulation of lipolysis in adipocytes, adipocytokine & cGMP-PKG signalling pathways, arachidonic acid metabolism and bile secretion. The results were validated by molecular docking which showed that RXRA, STAT3 and ESR1 strongly interact with their ligands alpha gurjunene, beta-sitosterol, piperlongumine, genistein and E-beta carotene, respectively.

**Conclusion:** Hence, the multiple target and multiple pathway approach of Trikatu can be further explored in pharmacokinetics / Pharmacodynamics studies.

## Introduction

Metabolic disorders have increased in prevalence over the last few decades(1). Globally, more than one billion young adults have diabetes, obesity, and dyslipidaemia which is associated with an increased risk of cardiovascular events like hypertension, Coronary Artery Disease (CAD), cerebrovascular stroke and increased mortality (2, 3). Metabolic disorders, including obesity, type 2 diabetes mellitus (T2DM), and nonalcoholic fatty liver disease (NAFLD), pose a considerable socioeconomic burden across the world (4). Current treatment of metabolic disorders includes LDL-cholesterol-lowering therapies, statins, omega-3 fatty acids, thiazolidinediones, renin-angiotensin system blockers, and peroxisome proliferator-activated receptor agonists (5, 6). However, these medications have a high cost, and are associated with various adverse effects, including weight gain, hypoglycemia, gastrointestinal problems, and hepatotoxicity (7, 8). Due to these limitations, patients are looking for the opportunity in complemenatry and alternative medicine for safe and cost effective. Moreover, there is a need to explore herb-based medications in complementary and alternative medicine (CAM) (5). Natural products and herbal formulations are cost-effective in treating non-communicable diseases (9–12).

In the Indian subcontinent, Ayurveda is one of the oldest and most popular complementary and alternative medicine systems (13). As per the classical texts of Ayurveda, herbs such as *Z. officinale* (ZO)*, P. nigrum* (PN)*, Piper longum* (PL)*, Phyllanthus emblica, Curcuma longa, Allium sativum, Cuminum cyminum* are commonly used to treat T2DM, obesity, skin disorders, dysentery, diarrhoea, flatulence, colic pain, chronic cough, liver diseases, bronchial asthma and cardiovascular disorders (14). *Trikatu* is an ayurvedic polyherbal formulation consisting of PL, PN and ZO in equal proportions (15, 16). Preclinical and clinical studies on *Trikatu* have shown to possess antiviral, carminative, hypolipidemic, hypoglycemic, antiemetic, anti-inflammatory, antihypothyroidism, antifungal and antibacterial properties (15,17,18). In clinical practice, it is prescribed to boost metabolism and manage chronic rhinitis/sinusitis, cough, asthma, diabetes, obesity, and dyslipidaemia by Ayurveda physicians (16, 19). In obese and hypercholesterolemic patients, research studies have shown that *Trikatu* reduces body weight, skinfold thickness and total cholesterol levels (20, 21). Despite its wide clinical applications, the therapeutic mechanism of action of *Trikatu* remains unknown.

Network pharmacology (NP) has been recently used in computational biology to identify the bioactive phytochemicals and potential targets of traditional medicinal plants, providing novel insights to elucidate their safety, efficacy and therapeutic mechanism (22, 23)(22, 24). It also provides a new framework for drug discovery by recognizing novel therapeutic targets, saving time and cost and being beneficial in Pharmacology (25). As this approach is based on the ‘one drug, multiple targets’ model, it is also helpful to understand the mechanism of action of polyherbal formulations (25).

Therefore, the present study aims to identify the bioactive phytochemicals and elucidate the mechanism of action of *Trikatu* in T2DM, obesity and dyslipidemia using the NP-based approach. The schematic workflow approach used in this study is shown in Figure 1.

**Figure 1.**
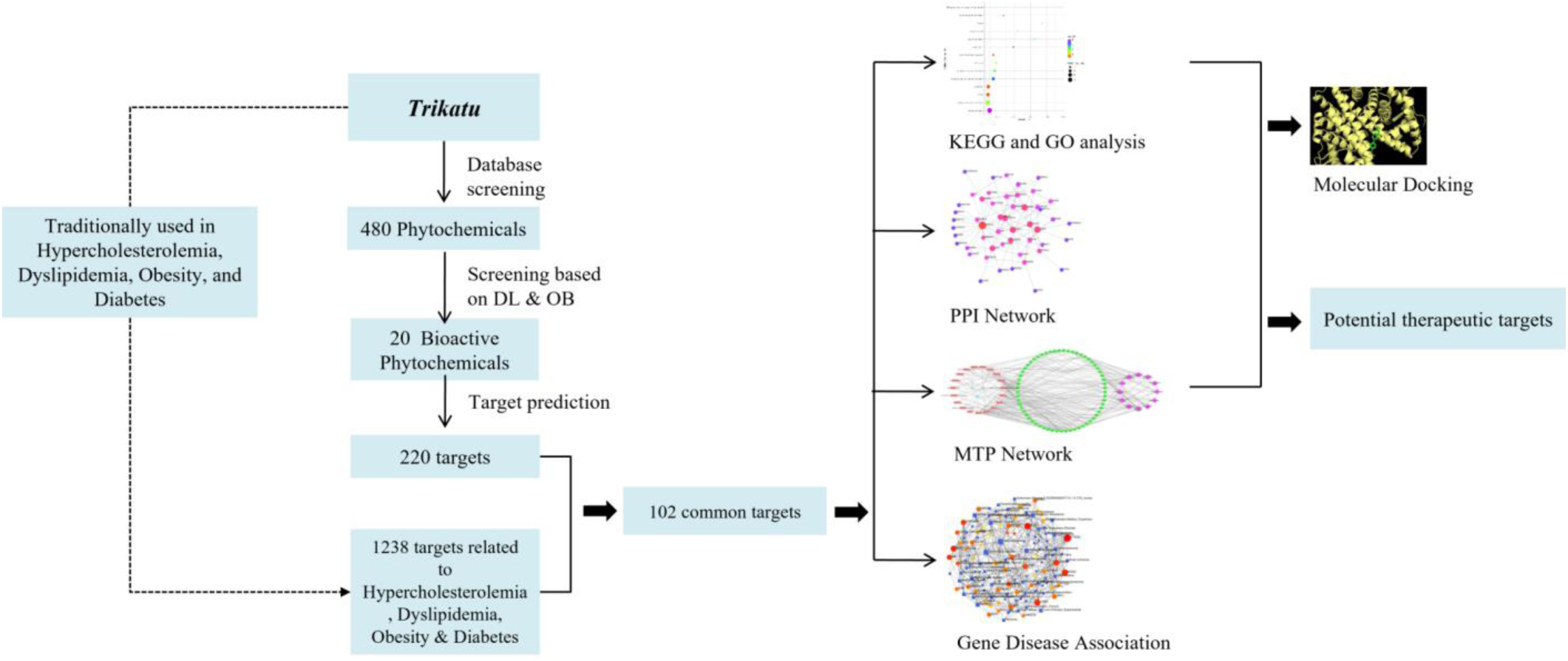
Schematic diagram of the workflow of network pharmacology of *Trikatu*.

## Materials and Methods

### Identification and screening of bioactive phytochemicals of *Trikatu*

The phytochemicals/metabolites of ZO, PL and PN were retrieved from PubChem (https://pubchem.ncbi.nlm.nih.gov/) (26), KNApSAcK (http://knapsackfamily.com/knapsack_core/top.php) (27), Indian Medicinal Plants, Phytochemistry and Therapeutics (IMPPAT, https://cb.imsc.res.in/imppat/) (28), and Chemical Entities of Biological Interest (ChEBI, https://www.ebi.ac.uk/chebi/) (29) databases, and screened based on drug-likeness (DL) and oral bioavailability (OB) using Molsoft (https://www.molsoft.com/) (30), and ADMETlab 2.0 (https://admet.scbdd.com/) (31). Phytochemicals with positive DL scores (32) and ≥ 30% OB were further analyzed based on ADMET (Absorption, Distribution, Metabolism, Excretion, and Toxicity) properties using admetSAR (http://lmmd.ecust.edu.cn/admetsar2) (33).

### Identification of target genes of *Trikatu* phytochemicals

The target genes of bioactive phytochemicals from *Trikatu* were identified using the similarity ensemble approach (SEA; http://sea.bkslab.org/) (34), TargetNet (http://targetnet.scbdd.com/) (35) and Swiss Target Prediction tool (http://www.swisstargetprediction.ch/) (36). Targets with Tc max ≥ 0.6 and probability ≥ 0.6 were selected for further analysis.

### Identification of targets related to T2DM and lipid metabolism

Genes related to obesity, dyslipidemia and T2DM were identified using databases, such as DisGeNET (0.08 gene-disease associations (GDA) score; https://www.disgenet.org/) (37), Online Mendelian Inheritance in Man (OMIM; http://www.omim.org/) (38), DrugBank (http://www.drugbank.ca/), Therapeutic Target Database (TTD; http://db.idrblab.net/ttd/) (39), and KEGG Disease (https://www.genome.jp/kegg/disease/) (40). The gene names were validated using the UniProt database (https://www.uniprot.org/) (41). The overlapping targets between *Trikatu* phytochemicals and genes related to metabolic disorders were selected using Venny v2.1.0 (https://bioinfogp.cnb.csic.es/tools/venny/) (42). The overlapping targets were further utilized to construct the PPI network and enrichment analysis.

### PPI network construction

PPI network was constructed using overlapping gene targets by NetworkAnalyst 3.0 tool (http://www.networkanalyst.ca/) (43). First, a zero-order network was created based on information from databases such as IntAct, BIND, MINT, BioGRID, and DIP which are integrated into InnateDB (44). The network was then imported into Cytoscape v3.8.2 (45) to analyze its properties using the inbuilt Network Analyser tool 3.0.

### Gene ontology (GO) and KEGG pathway enrichment analysis

GO enrichment analysis and KEGG pathway annotation were carried out using DAVID (https://david.ncifcrf.gov/) (46). GO terms (biological function, molecular function and cellular component) and KEGG pathways with *P* ≤ 0.05 (FDR and Benjamini-Hochberg) were selected. The rich factor was calculated to analyze the number of significant genes involved in the particular pathway. The enrichment analysis has been shown in a bubble plot made using the ggplot2 package in R software (47, 48).

### Construction of bioactive phytochemicals–target–pathway (BP-T-P) network

For a deeper understanding of the mechanistic action of *Trikatu,* a bioactive phytochemical – target–pathway (BP-T-P) network was constructed using cytoscape v3.8.2. The SIF file containing *Trikatu* bioactive phytochemicals/metabolites, their target genes, and the related KEGG pathways were all imported into Cytoscape v3.8.2 and a network was constructed in a degree sorted circle layout. The network properties were analyzed by the Network Analyser tool of Cytoscape v3.8.2.

### Molecular docking analysis

Molecular docking was conducted using hub proteins with degrees ≥ 10 in the PPI network. The ligand information of the proteins was obtained from the BP-T-P network. The structure of proteins and ligands were obtained from Protein Data Bank (PDB; www.rcsb.org) (49) and PubChem, respectively. The binding site of the protein molecule was determined by CASTp (http://sts.bioe.uic.edu/castp/index.html?1ycs) (50). The proteins and the ligands were prepared before docking by removing water molecules, adding polar hydrogen, and Kollman charges (51). The grid map and size of the protein molecule were calculated to access the binding area for the ligand. Subsequently, the protein and ligand were subjected to docking using Autodock Vina v1.1.2 (52), which uses Broyden-Fletcher-Goldfarb-Shanno (BFGS) algorithm. The highest negative binding affinities conformer were selected and visualized using PyMOL v2.5. The interaction between ligand and protein complex was analysed by protein-ligand interaction profiler (https://plip-tool.biotec.tu-dresden.de/plip-web/plip/index) (53) and LigPlot+ v.2.2 (https://www.ebi.ac.uk/thornton-srv/software/LigPlus/) (54).

### Gene-disease association network construction

The association of target genes with other diseases was analyzed via the ‘gene-disease associations’ network mapping tool available on the Network Analyst 3.0 platform. This tool uses the literature curated gene-disease association data using the DisGeNET database (https://www.disgenet.org/) (37).

## Results

### Identification of bioactive phytochemicals of *Trikatu*

A total of 480 metabolites were identified for PL, PN and ZO (Supplementary File 1). Further, out of 480 phytochemicals, 20 bioactive phytochemicals were filtered on the basis of OB (≥ 30%) and positive DL score. Beta-sitosterol (0.78), phytosterol (0.78), ascorbic acid (0.74), paroxetine (0.78), dihyroferuperine (0.74), and E-beta carotene (0.64) had higher drug-likeness score (Table 1). Camphor (0.731) and paroxetine (0.683) were found to have higher oral bioavailability scores (Table 1). ADMET profiles, physiochemical and drug-likeness properties of bioactive phytochemicals have been shown in in the form of heatmap in Supplementary File 2.

**Table 1.**
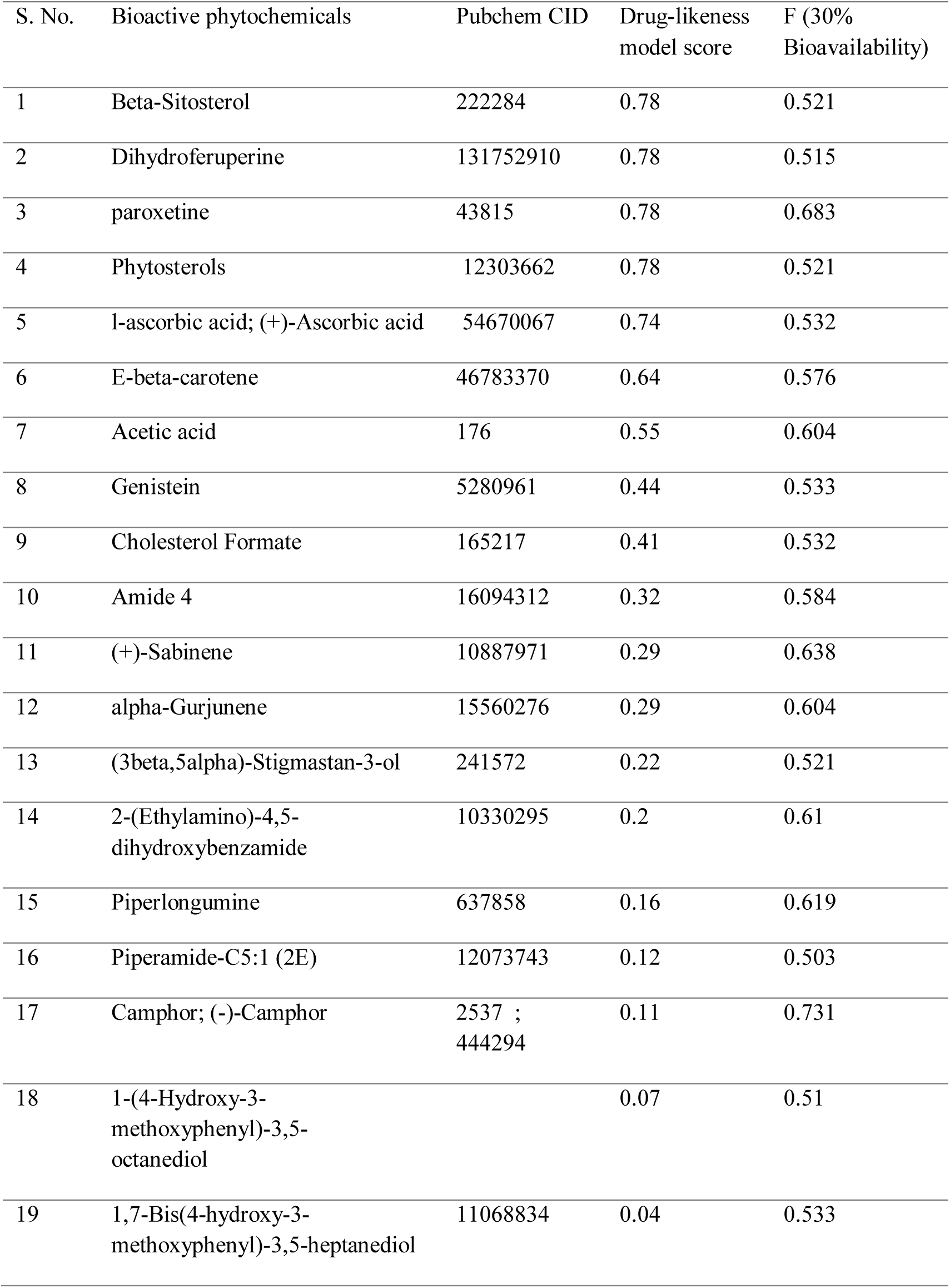
Bioactive phytochemicals of *Trikatu*

| S. No. | Bioactive phytochemicals | Pubchem CID | Drug-likeness model score | F (30% Bioavailability) |
| --- | --- | --- | --- | --- |
| 1 | Beta-Sitosterol | 222284 | 0.78 | 0.521 |
| 2 | Dihydroferuperine | 131752910 | 0.78 | 0.515 |
| 3 | paroxetine | 43815 | 0.78 | 0.683 |
| 4 | Phytosterols | 12303662 | 0.78 | 0.521 |
| 5 | l-ascorbic acid; (+)-Ascorbic acid | 54670067 | 0.74 | 0.532 |
| 6 | E-beta-carotene | 46783370 | 0.64 | 0.576 |
| 7 | Acetic acid | 176 | 0.55 | 0.604 |
| 8 | Genistein | 5280961 | 0.44 | 0.533 |
| 9 | Cholesterol Formate | 165217 | 0.41 | 0.532 |
| 10 | Amide 4 | 16094312 | 0.32 | 0.584 |
| 11 | (+)-Sabinene | 10887971 | 0.29 | 0.638 |
| 12 | alpha-Gurjunene | 15560276 | 0.29 | 0.604 |
| 13 | (3beta,5alpha)-Stigmastan-3-ol | 241572 | 0.22 | 0.521 |
| 14 | 2-(Ethylamino)-4,5-dihydroxybenzamide | 10330295 | 0.2 | 0.61 |
| 15 | Piperlongumine | 637858 | 0.16 | 0.619 |
| 16 | Piperamide-C5:1 (2E) | 12073743 | 0.12 | 0.503 |
| 17 | Camphor; (-)-Camphor | 2537 ;<br>444294 | 0.11 | 0.731 |
| 18 | 1-(4-Hydroxy-3-methoxyphenyl)-3,5-octanediol |  | 0.07 | 0.51 |
| 19 | 1,7-Bis(4-hydroxy-3-methoxyphenyl)-3,5-heptanediol | 11068834 | 0.04 | 0.533 |

### Target identification of *Trikatu* bioactive phytochemicals

A total of 871 targets of bioactive phytochemicals having Tc max ≥ 0.6 and probability ≥ 0.6 were identified (Supplementary File 1). Further, 220 protein targets were selected after the removal of duplicate entries. On the other hand, 1238 genes related to obesity, hypercholesteremia, T2DM and dyslipidaemia were identified (Supplementary File 1). Finally, 102 overlapping genes between targets of phytochemicals and metabolic disorders were selected and used to construct the PPI network.

### PPI network construction

PPI network constructed using overlapping targets consist of 51 nodes and 100 edges with clustering coefficient 0.119 and network density 0.078. The degree and betweenness of the nodes ranged from 22 to 1 and 593.93 to 1, respectively (Figure 2; Table 2). Results showed that the nodes with a high degree and high betweenness were hub genes with size generally bigger than other nodes. Target genes such as HNF4A, RXRA, ESR1, SIRT1, and STAT3 showing degree ≥ 10 were considered hub genes (Table 2).

**Figure 2.**
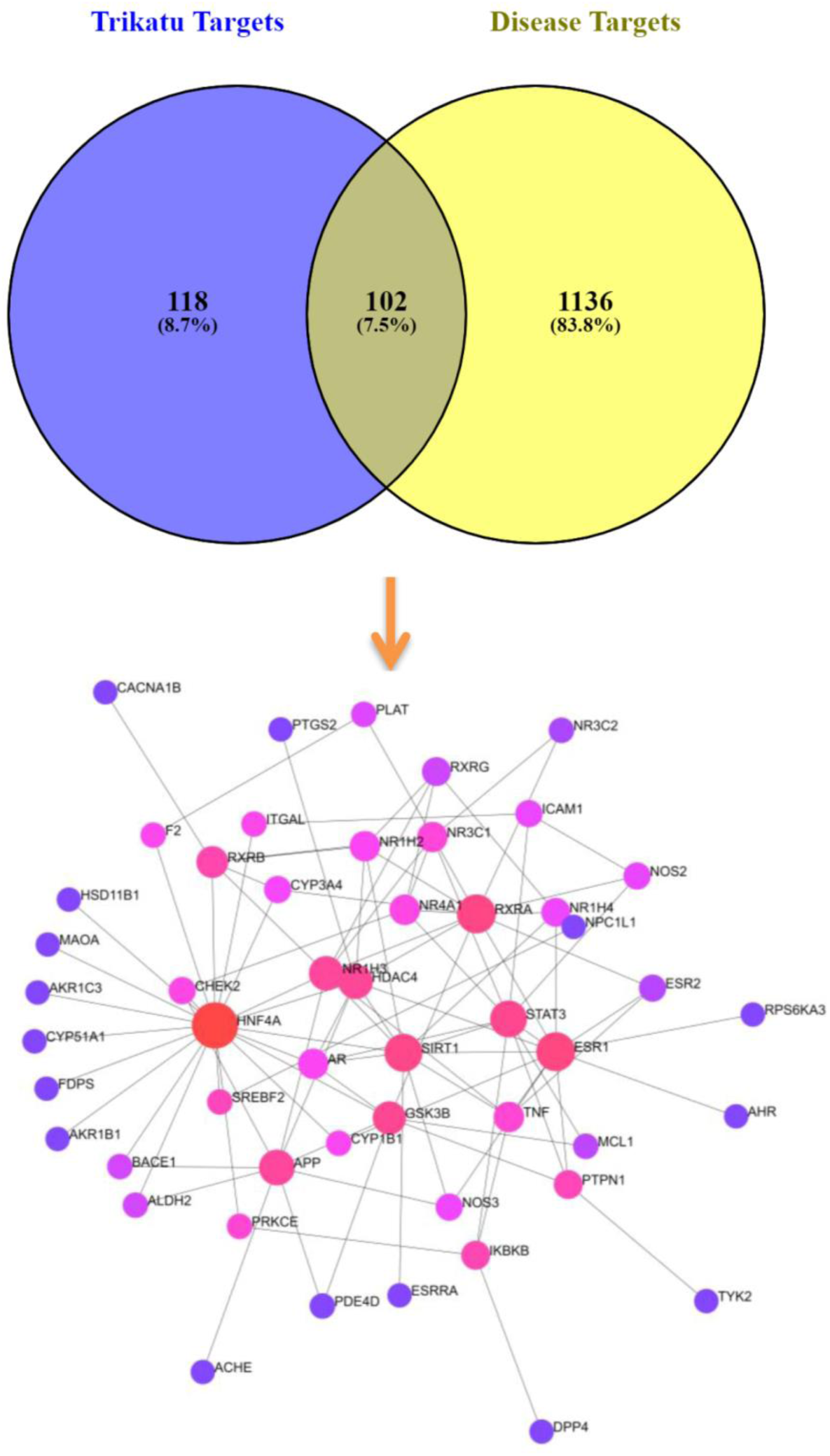
PPI interaction network of overlapping genes between *Trikatu* targets and disease targets such as obesity, hypercholesterolemia, dyslipidaemia and obesity. The node size and colour gradient (The gradient colour: red→ pink → purple; and size: large to small) represent the high to the low degree of the nodes within the network.

**Table 2.** Topological properties of Protein-protein interaction network

| Protein | Average Shortest Path Length | Betweenness Centrality | Closeness Centrality | Clustering Coefficient | Degree | Eccentricity | Topological Coefficient |
| --- | --- | --- | --- | --- | --- | --- | --- |
| SIRT1 | 1.94 | 0.121 | 0.515 | 0.218 | 11 | 3 | 0.180 |
| ESR1 | 2.18 | 0.138 | 0.459 | 0.152 | 12 | 4 | 0.182 |
| HDAC4 | 2.02 | 0.085 | 0.495 | 0.250 | 8 | 4 | 0.231 |
| HNF4A | 1.74 | 0.469 | 0.575 | 0.033 | 21 | 3 | 0.097 |
| NR1H3 | 2.04 | 0.081 | 0.490 | 0.143 | 8 | 4 | 0.230 |
| STAT3 | 2.2 | 0.104 | 0.455 | 0.111 | 10 | 4 | 0.173 |
| NOS3 | 2.52 | 0.007 | 0.397 | 0.333 | 3 | 4 | 0.449 |
| AR | 2.12 | 0.012 | 0.472 | 0.500 | 5 | 3 | 0.292 |
| NR1H2 | 2.46 | 0.012 | 0.407 | 0.200 | 5 | 4 | 0.339 |
| ESRRA | 2.92 | 0.000 | 0.342 | 0.000 | 1 | 4 | 0.000 |
| NR1H4 | 2.52 | 0.007 | 0.397 | 0.333 | 4 | 4 | 0.413 |
| TNF | 2.44 | 0.026 | 0.410 | 0.300 | 5 | 4 | 0.322 |
| APP | 2.42 | 0.078 | 0.413 | 0.028 | 9 | 4 | 0.201 |
| PDE4D | 2.86 | 0.000 | 0.350 | 1.000 | 2 | 5 | 0.571 |
| GSK3B | 2.12 | 0.095 | 0.472 | 0.095 | 7 | 4 | 0.212 |
| CHEK2 | 2.42 | 0.017 | 0.413 | 0.000 | 3 | 4 | 0.395 |
| ALDH2 | 2.6 | 0.004 | 0.385 | 0.000 | 2 | 4 | 0.609 |
| ACHE | 3.4 | 0.000 | 0.294 | 0.000 | 1 | 5 | 0.000 |
| BACE1 | 2.6 | 0.004 | 0.385 | 0.000 | 2 | 4 | 0.609 |
| NOS2 | 2.78 | 0.006 | 0.360 | 0.333 | 3 | 5 | 0.400 |
| RXRA | 2.16 | 0.125 | 0.463 | 0.061 | 12 | 4 | 0.183 |
| ICAM1 | 2.82 | 0.007 | 0.355 | 0.333 | 3 | 5 | 0.389 |
| AHR | 3.16 | 0.000 | 0.316 | 0.000 | 1 | 5 | 0.000 |
| RPS6KA3 | 3.16 | 0.000 | 0.316 | 0.000 | 1 | 5 | 0.000 |
| ESR2 | 2.8 | 0.002 | 0.357 | 0.667 | 3 | 5 | 0.460 |
| PTPN1 | 2.48 | 0.044 | 0.403 | 0.167 | 4 | 4 | 0.318 |
| CYP1B1 | 2.38 | 0.012 | 0.420 | 0.000 | 2 | 4 | 0.534 |
| PTGS2 | 3 | 0.000 | 0.333 | 0.000 | 1 | 5 | 0.000 |
| ITGAL | 2.62 | 0.015 | 0.382 | 0.000 | 2 | 4 | 0.500 |
| CYP3A4 | 2.36 | 0.009 | 0.424 | 0.333 | 3 | 4 | 0.409 |
| SREBF2 | 2.68 | 0.040 | 0.373 | 0.000 | 2 | 4 | 0.500 |
| PRKCE | 2.56 | 0.026 | 0.391 | 0.000 | 2 | 4 | 0.500 |
| AKR1B1 | 2.72 | 0.000 | 0.368 | 0.000 | 1 | 4 | 0.000 |
| FDPS | 2.72 | 0.000 | 0.368 | 0.000 | 1 | 4 | 0.000 |
| F2 | 2.6 | 0.013 | 0.385 | 0.000 | 2 | 4 | 0.500 |
| CYP51A1 | 2.72 | 0.000 | 0.368 | 0.000 | 1 | 4 | 0.000 |
| MAOA | 2.72 | 0.000 | 0.368 | 0.000 | 1 | 4 | 0.000 |
| AKR1C3 | 2.72 | 0.000 | 0.368 | 0.000 | 1 | 4 | 0.000 |
| RXRB | 2.34 | 0.054 | 0.427 | 0.200 | 6 | 4 | 0.236 |
| HSD11B1 | 2.72 | 0.000 | 0.368 | 0.000 | 1 | 4 | 0.000 |
| IKBKB | 2.72 | 0.048 | 0.368 | 0.000 | 4 | 4 | 0.292 |
| DPP4 | 3.7 | 0.000 | 0.270 | 0.000 | 1 | 5 | 0.000 |
| NR3C2 | 3 | 0.001 | 0.333 | 0.000 | 2 | 5 | 0.577 |
| NR3C1 | 2.5 | 0.022 | 0.400 | 0.200 | 5 | 4 | 0.286 |
| RXRG | 2.68 | 0.003 | 0.373 | 0.000 | 4 | 4 | 0.409 |
| NR4A1 | 2.42 | 0.018 | 0.413 | 0.100 | 5 | 4 | 0.282 |
| MCL1 | 2.72 | 0.002 | 0.368 | 0.000 | 2 | 4 | 0.577 |
| PLAT | 2.84 | 0.005 | 0.352 | 0.000 | 2 | 5 | 0.500 |
| NPC1L1 | 3.66 | 0.000 | 0.273 | 0.000 | 1 | 5 | 0.000 |
| TYK2 | 3.46 | 0.000 | 0.289 | 0.000 | 1 | 5 | 0.000 |
| CACNA1B | 3.32 | 0.000 | 0.301 | 0.000 | 1 | 5 | 0.000 |

### GO and KEGG pathway enrichment analysis

To explore the molecular mechanism of *Trikatu* in metabolic disorders, significant GO terms with *P* ≤ 0.05 were selected (Supplementary File 3). The biological GO terms and pathways were found to be mainly involved in lipid and glucose homeostasis. The significant biological processes were steroid hormone-mediated signalling pathway, regulation of insulin secretion, adenylate cyclase-activating adrenergic receptor signalling pathway, regulation of cholesterol homeostasis, glucose homeostasis, retinoic acid receptor signalling pathway, activation of cholesterol biosynthetic process, negative regulation of lipid catabolic process etc. (Figure 3A). The target proteins were found to be involved in various molecular functions such as steroid hormone receptor activity, steroid binding, receptor binding, epinephrine binding, transcriptional activator activity, and iron ion binding retinoid X receptor binding etc. were found to be significant (Figure 3B). The results also showed that the target proteins are associated with different cellular components like plasma membrane, endoplasmic reticulum membrane, receptor complex, cell surface, cytosol, nucleoplasm etc. (Figure 3C).

**Figure 3.**
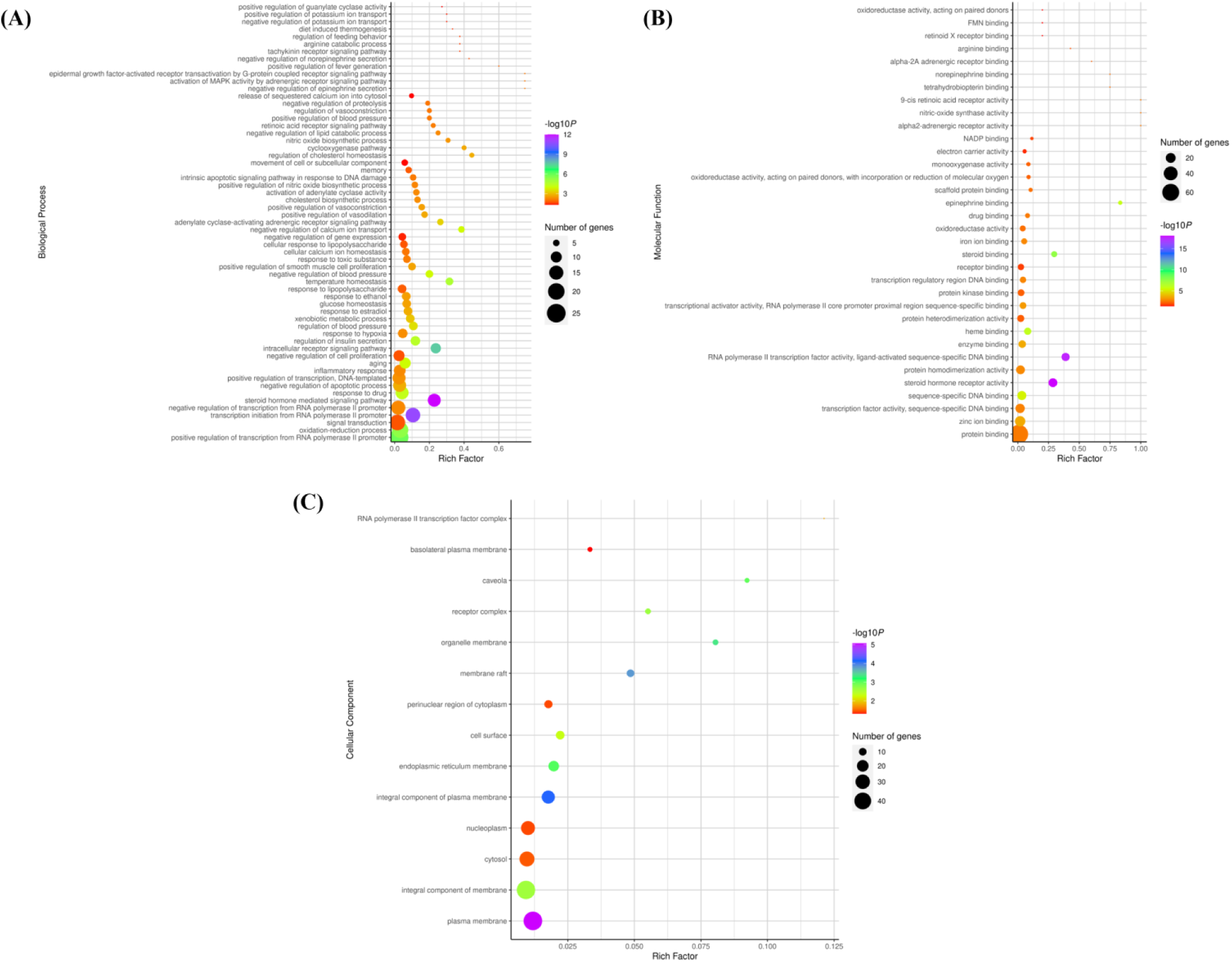
Gene Ontology enrichment analysis. (A) Biological function (B) Molecular function C) Cellular Component. The log *P* values and rich factor was calculated to show the number of significant genes involved in that pathway term. Rich factor is defined as the ratio of number of genes involved in a given pathway term to the total number of genes annotated in that pathway term. Higher the rich factor and greater will be the gene enrichment in the given pathway.

In our study, KEGG pathways with *P* ≤ 0.05 were found to be neuroactive ligand-receptor interaction, insulin resistance, calcium signalling pathway, steroid hormone biosynthesis, serotonergic synapse, regulation of lipolysis in adipocytes, adipocytokine signalling pathway, arginine and proline metabolism, thyroid hormone signalling pathway, small cell lung cancer, cGMP-PKG signalling pathway, arachidonic acid metabolism, and bile secretion (Figure 4). These pathways are mainly related to the etiology of metabolic diseases regulated by *Trikatu*.

**Figure 4.**
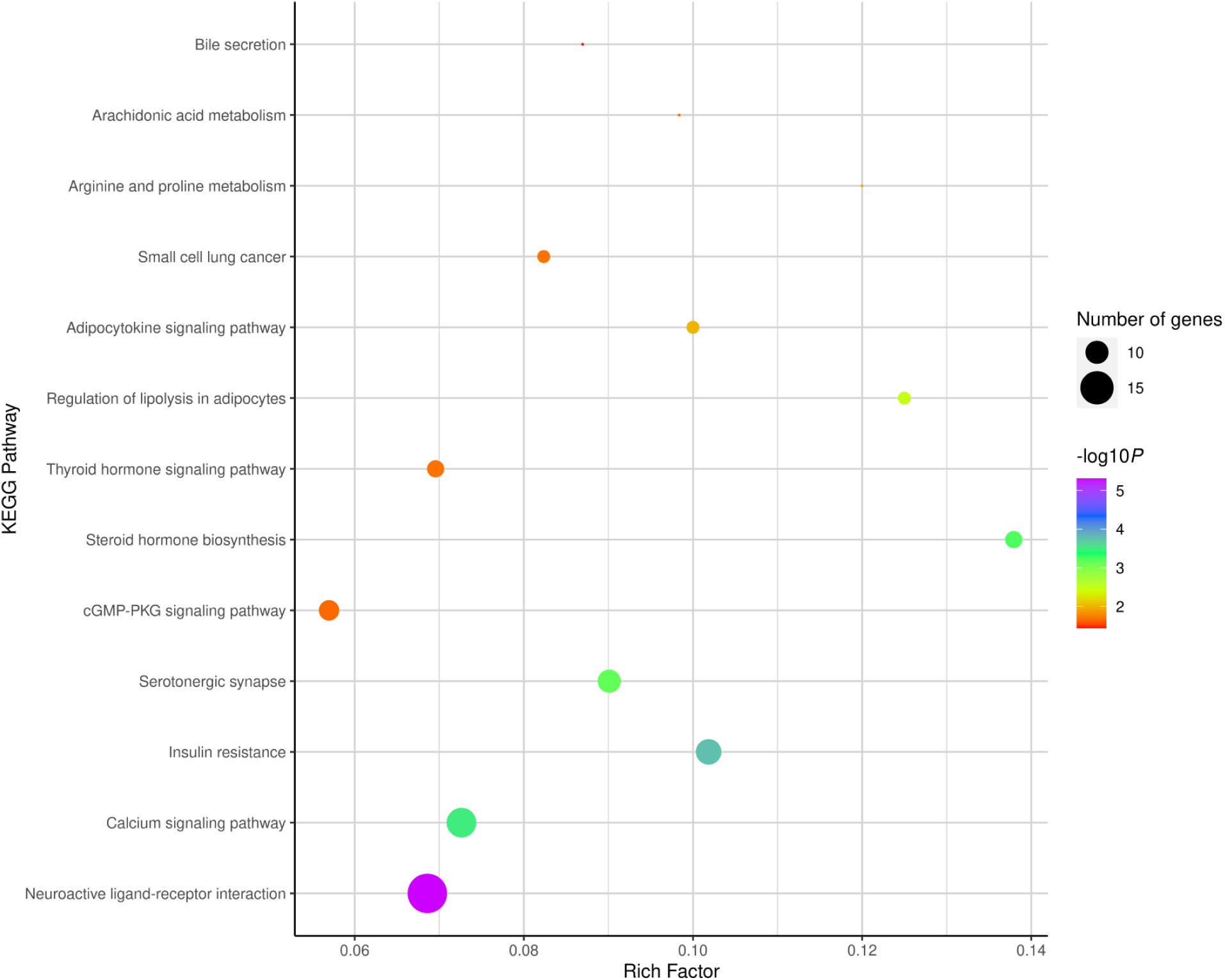
KEGG pathway using ggplot2 with R package. The *log P* values and rich factor was calculated to show the number of significant genes involved in that pathway term. Rich factor is defined as the ratio of number of genes involved in given pathway term to the total number of genes annotated in that pathway term. Higher the rich factor and greater will be the gene enrichment in the given pathway.

### Bioactive phytochemical-Target-Pathway (BP-T-P) network

The bioactive phytochemicals-target-pathway network consisted of 95 nodes and 305 edges with a network density of 0.068 (Figure 5). In this network, phytochemicals such as acetic acid, genistein, camphor, paroxetine, phytosterols, alpha-gurjunene, 1,7-Bis (4-hydroxy-3-methoxyphenyl)-3,5-heptanediol, beta-Sitosterol, and sabinene possess degree ≥ 6 (Supplementary File 4). The network showed that phytochemicals are associated with multiple target proteins regulating various pathways (Table 3). Some of the important pathways related to obesity, dyslipidaemia, and T2DM were insulin resistance, regulation of lipolysis in adipocytes, arachidonic acid metabolism, adipocyte signalling pathway, and bile secretion.

**Figure 5.**
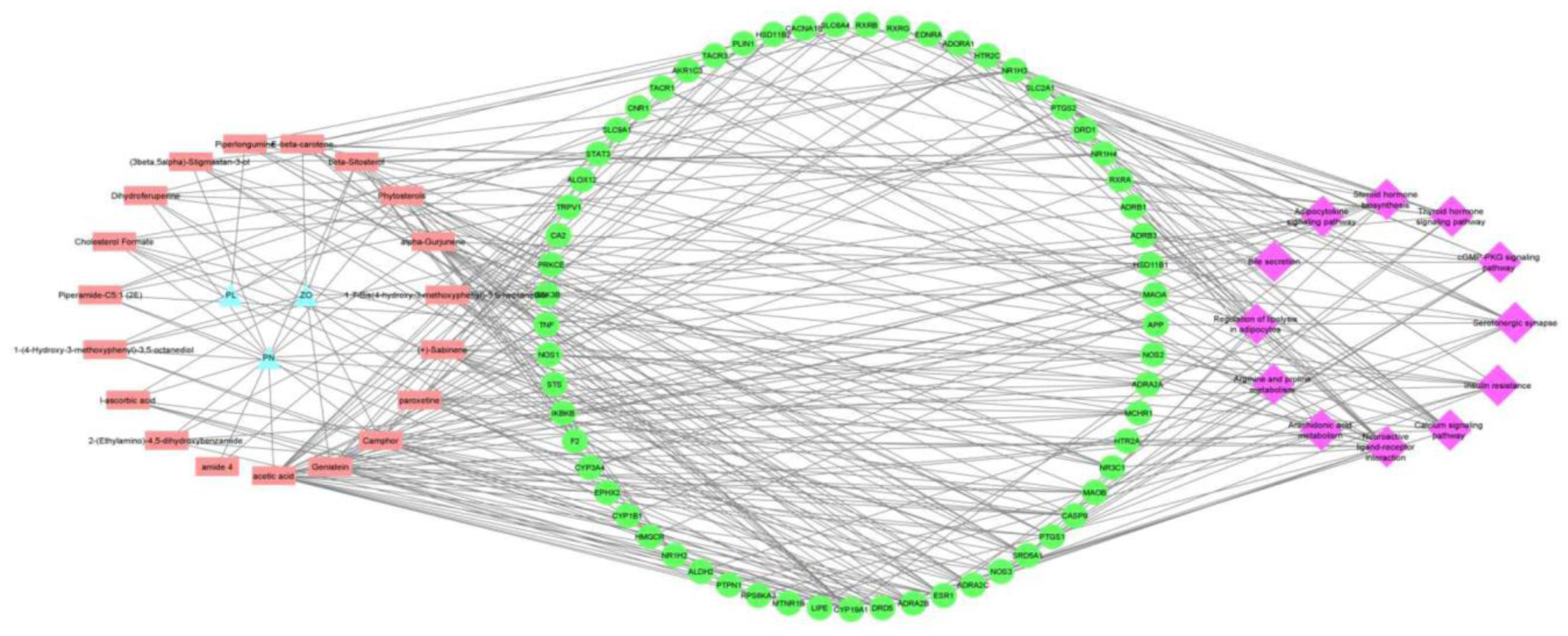
Bioactive phytochemical-Target-Pathway interaction network. The network was constructed in the form of a degree sorted network using Cytoscape v3.8.2. Brick red square box represents metabolites, pink diamond represents pathway and light green circle represents protein involved in diabetes, dyslipidemia, and obesity.

**Table 3.**
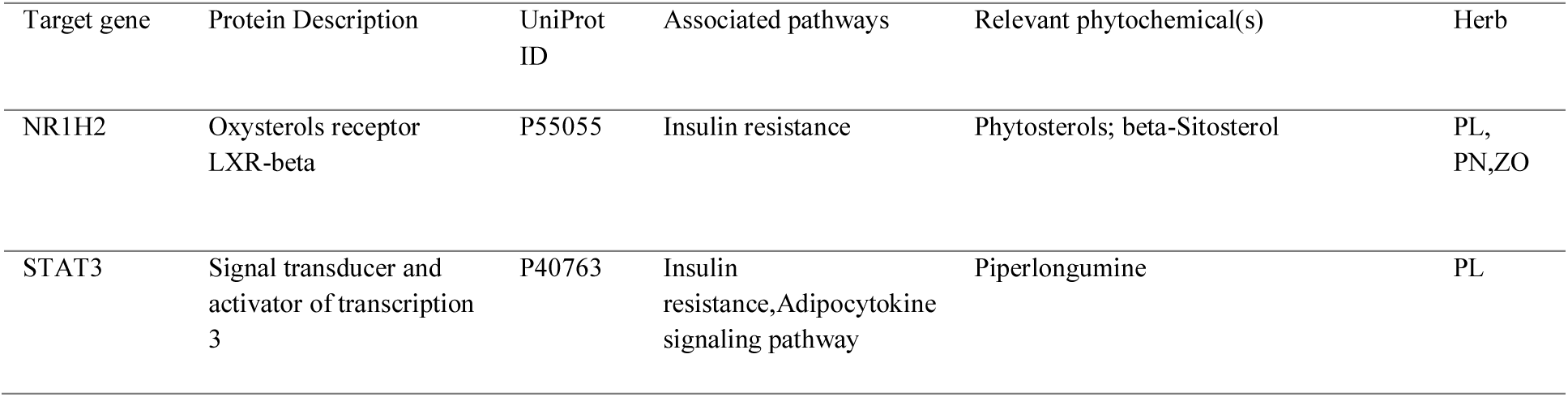
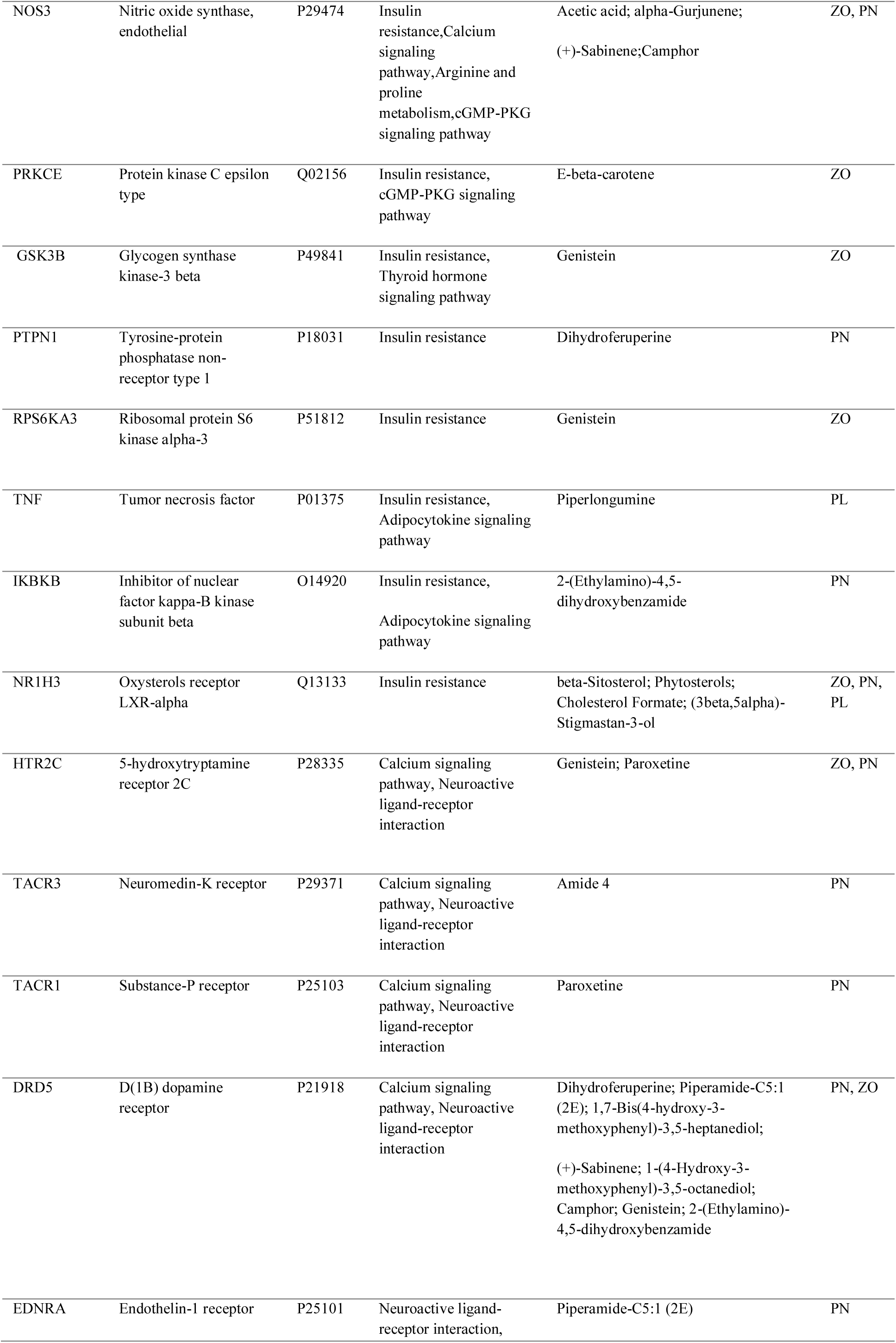
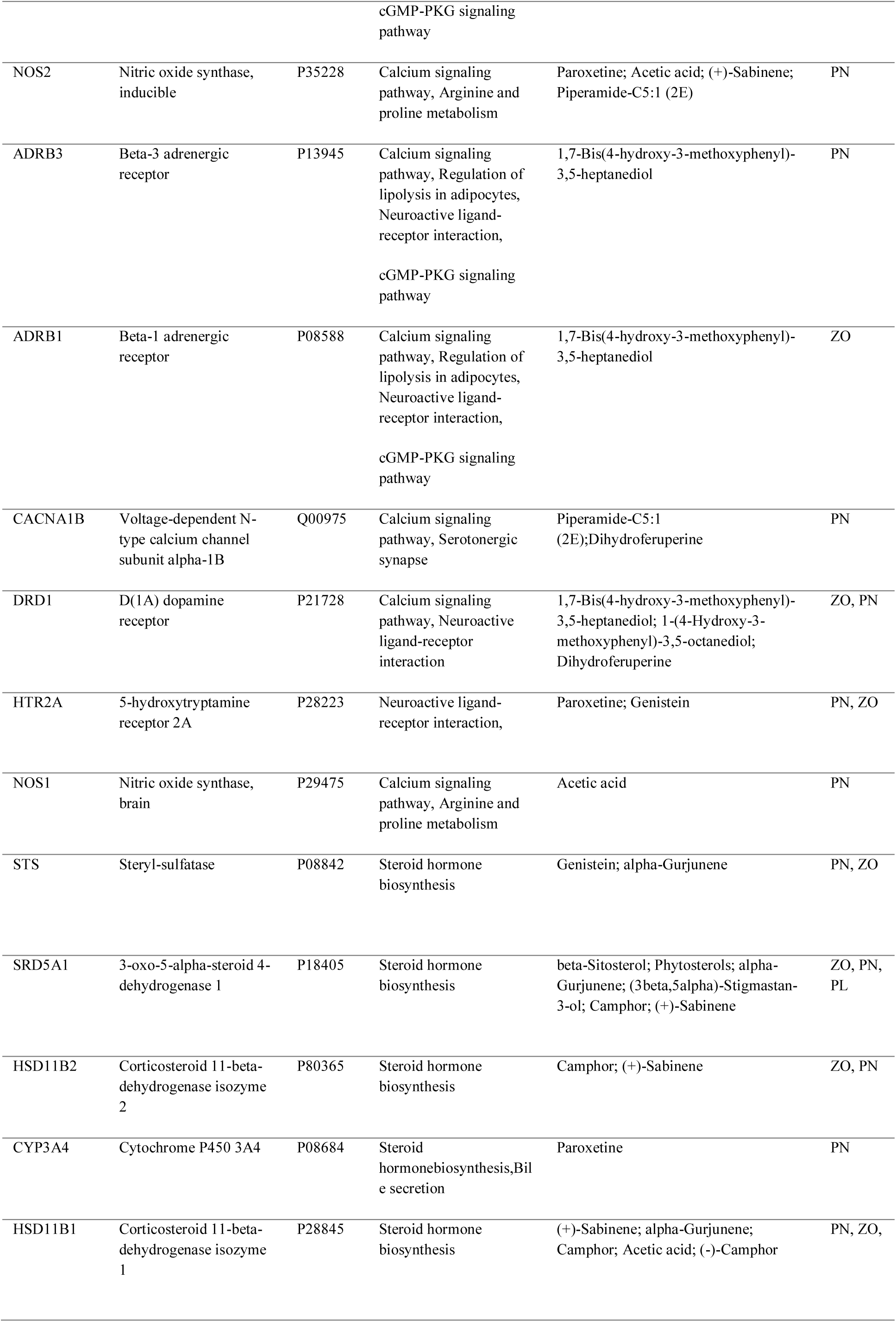
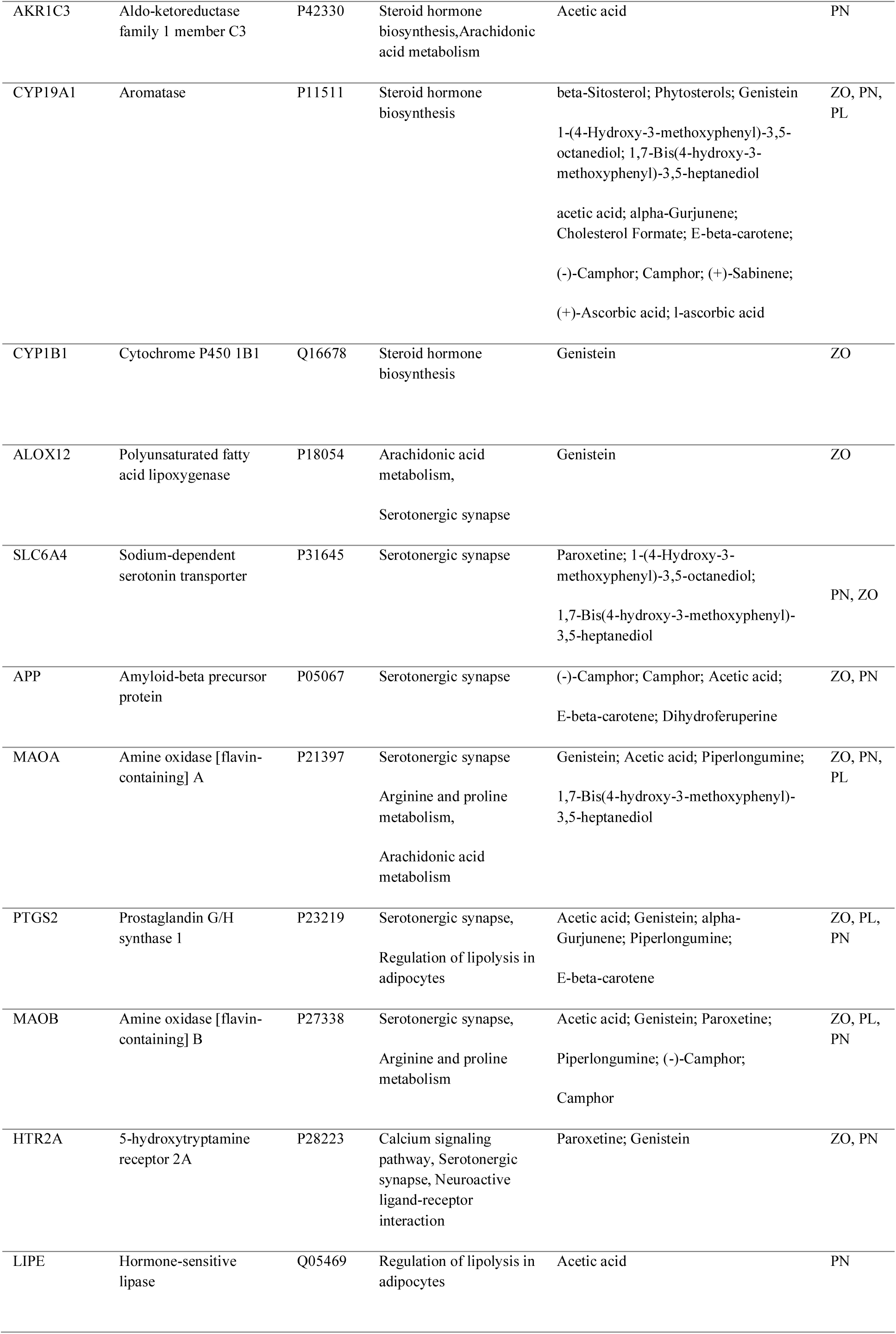
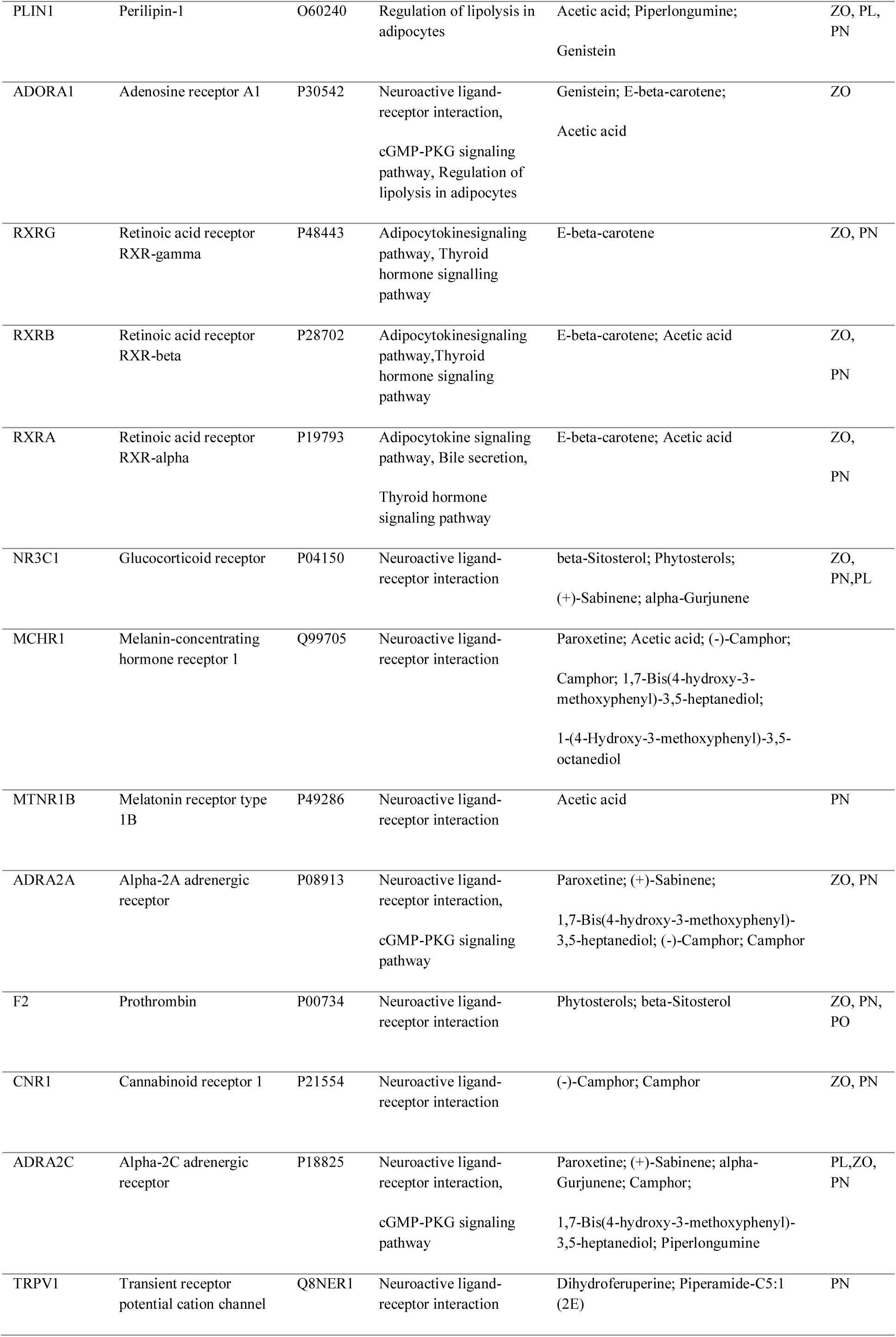
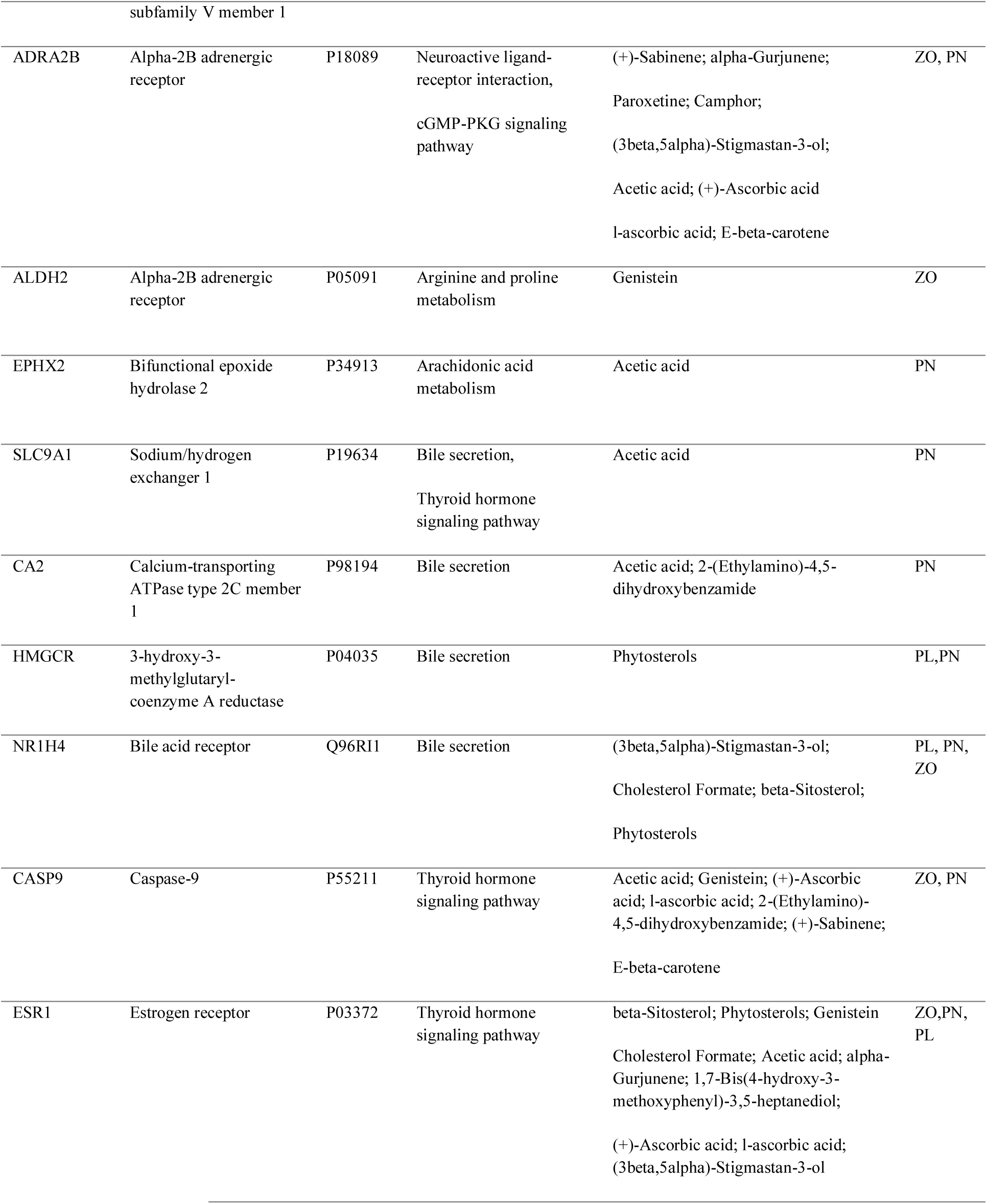
Targets of *Trikatu* related to metabolic disorders

### Molecular docking of protein and phytochemicals

Molecular docking was used to further confirm the interaction of phytochemicals and target proteins in the BP-T-P network. RXRA, STAT-3, and ESR1 proteins having degrees > 10 were selected for protein-ligand docking. A higher negative binding affinity value corresponds to strong binding between the ligand (i.e., metabolite) and the target protein. Results showed that emtabolites such as genistein, alpha gurjnene, beta-sitosterol, and acetic acid were found to interact with ESR1 protein, while acetic acid and E-beta carotene interacted with RXRA. ESR1 showed a higher affinity to genistein (−8.2 kcal/mol) and alpha gurjnene (−7.2 kcal/mol), and beta-sitosterol (−6.2 kcal/mol) (Figure 6, Table 3). Similarly, RXRA binds more strongly with E-beta carotene (−6.7 kcal/mol) compared to acetic acid (−3.4 kcal/mol) (Table 4). STAT3’s binding affinity to piperlongumine was determined to be - 6.3 kcal/mol. The detailed information regarding various interactions between protein and ligands has been shown in Table 4 and Supplementary File 5.

**Figure 6.**
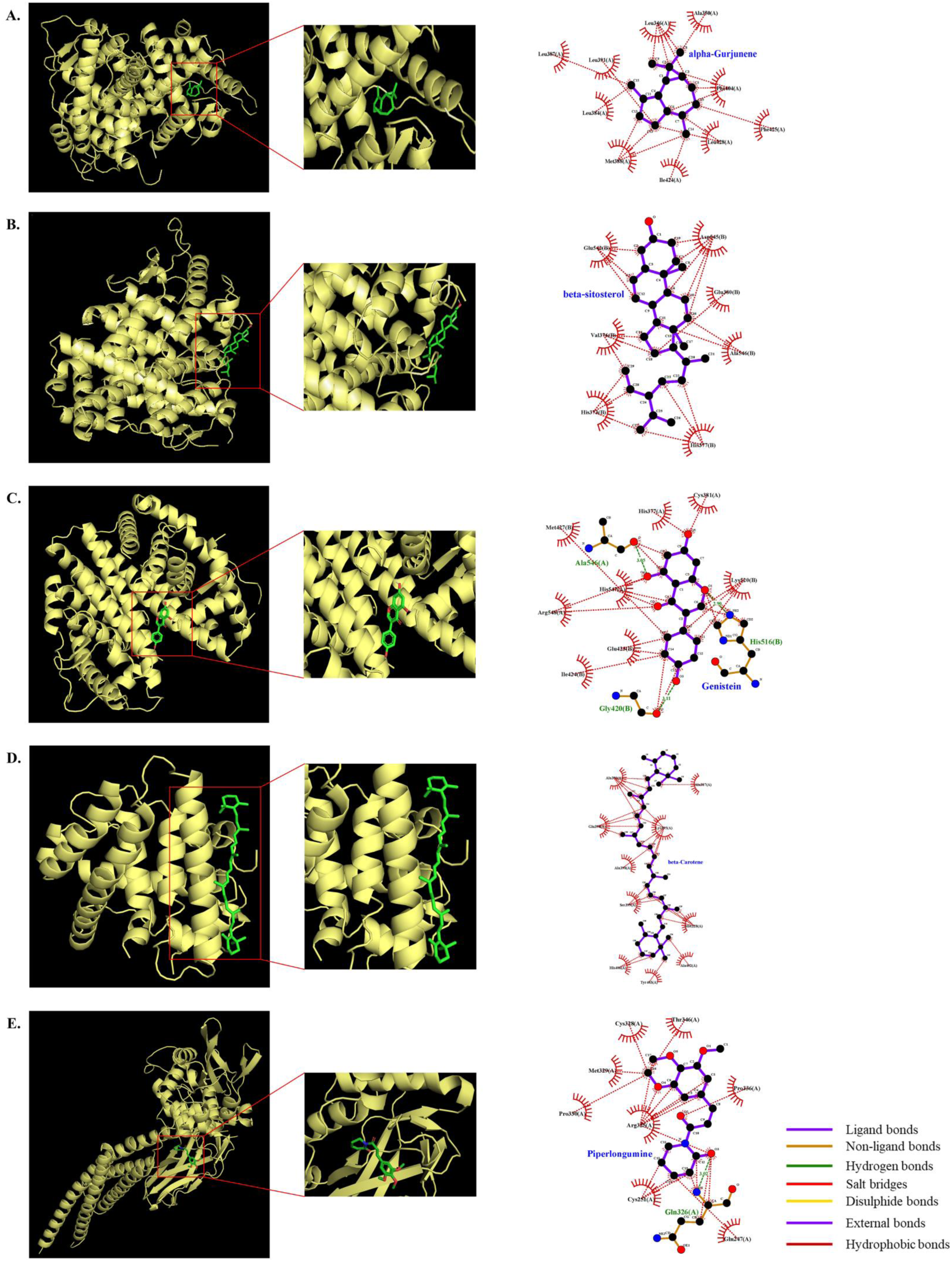
Docking of ESR with Genistein, beta-sitosterol, alpha gurjunene, RXRA with beta carotene, STAT3 with piperlongumine.

**Table 4:** Molecular docking of *Trikatu* targets with their ligands

| Protein | Ligand | Binding Affinity<br>(kcal/mol) | Amino acid Residues | Type of Interaction |
| --- | --- | --- | --- | --- |
| ESR1<br>(PDB ID:<br>5DTV) | Genistein<br>(PubChem ID:<br>5280961) | -7.2 | Glu423, Ile424, Lys520<br>Gly420, Ala546<br>His547 | Hydrophobic interactions<br>Hydrogen bonds<br>Pie interactions |
|  | Acetic acid<br>(PubChem ID:<br>176) | -3.0 | Leu387 | Hydrophobic interactions |
|  | beta-Sitosterol<br>(PubChem ID:<br>222284) | -6.2 | His373, Val376, His377,<br>Glu380, Glu542, Asp545 | Hydrophobic interactions |
|  | alpha-Gurjunene<br>(PubChem ID:<br>15560276) | -8.2 | Leu346, Ala350,<br>Leu384, Leu387,<br>Ile424, Phe425 | Hydrophobic interactions |
|  | Ascorbic acid<br>(Pubchem ID:<br><u>54670067</u>) | -4.4 | Ala430, Ser433, Tyr459,<br>Phe461, Lys472 | Hydrogen bond and salt bridges |
|  | Cholesterol<br>Formate<br>(PubChem ID:<br>165217) | -5.7 | His373, Val376, His373,<br>His377, Glu380, Glu542,<br>Ala546, Lys688 | Hydrophobic interaction |
|  | (3beta,5alpha)-<br>Stigmastan-3-ol<br>(PubChem ID:<br><u>241572</u> ) | -5.3 | Gln441, Ala493, Leu495,<br>Met438, Gln441, Glu444,<br>Asn439, Ala493, Leu495 | Hydrophobic interaction<br>Hydrogen Bond |
|  | 1,7-Bis(4-hydroxy-3-methoxyphenyl)<br>-3,5-heptanediol<br>( PubChem ID:<br><u>11068834</u>) | -5.8 | Ile424, Lys520, Glu523,<br>Arg548, Leu544, His547,<br>Arg548, Tyr526, Lys520,<br>Met522, Arg548 | Hydrophobic Interaction, hydrogen<br>Bonding, pie stacking, pie cation<br>interaction |
| RXRA<br>(PDB ID: | Acetic Acid | -3.4 | Trp305, Arg302 | Hydrogen bonds |
| 7B88) | (PubChem ID: 176) |  |  | Salt bridges interactions |
|  | beta-carotene | -6.7 | Met228, Val232, Ala387, Ala391, Lys395, Ala402, His406 | Hydrophobic interactions |
|  | (PubChem ID: 5280489) |  |  |  |
| STAT3 | Piperlongumine | -6.3 | Gln247, Ala250, Gln326, | Hydrogen bond, Hydrophobic interaction |
| (PDB ID: 6NJS) | (PubChem ID: 637858) |  | Thr346, Cys251, Met329, Prol336, Arg325, Cys328 |  |

### Gene-disease association network

Gene-disease association was analyzed using Network Analyst 3.0. The significant associations with *P* ≤ 0.05 were considered significant (Figure 7). The degree and betweenness of the resultant diseases ranged from 27 to 2 and 1231 to 0.45, respectively (Supplementary File 4). The results showed that *Trikatu* could also be explored in various diseases such as schizophrenia, adenocarcinoma, depressive disorder, mammary neoplasms, prostatic neoplasms, hypertensive disease, liver cirrhosis, bipolar disorder, liver carcinoma, seizures, stomach neoplasms, atherosclerosis, inflammation, myocardial infarction, fever, etc.

**Figure 7.**
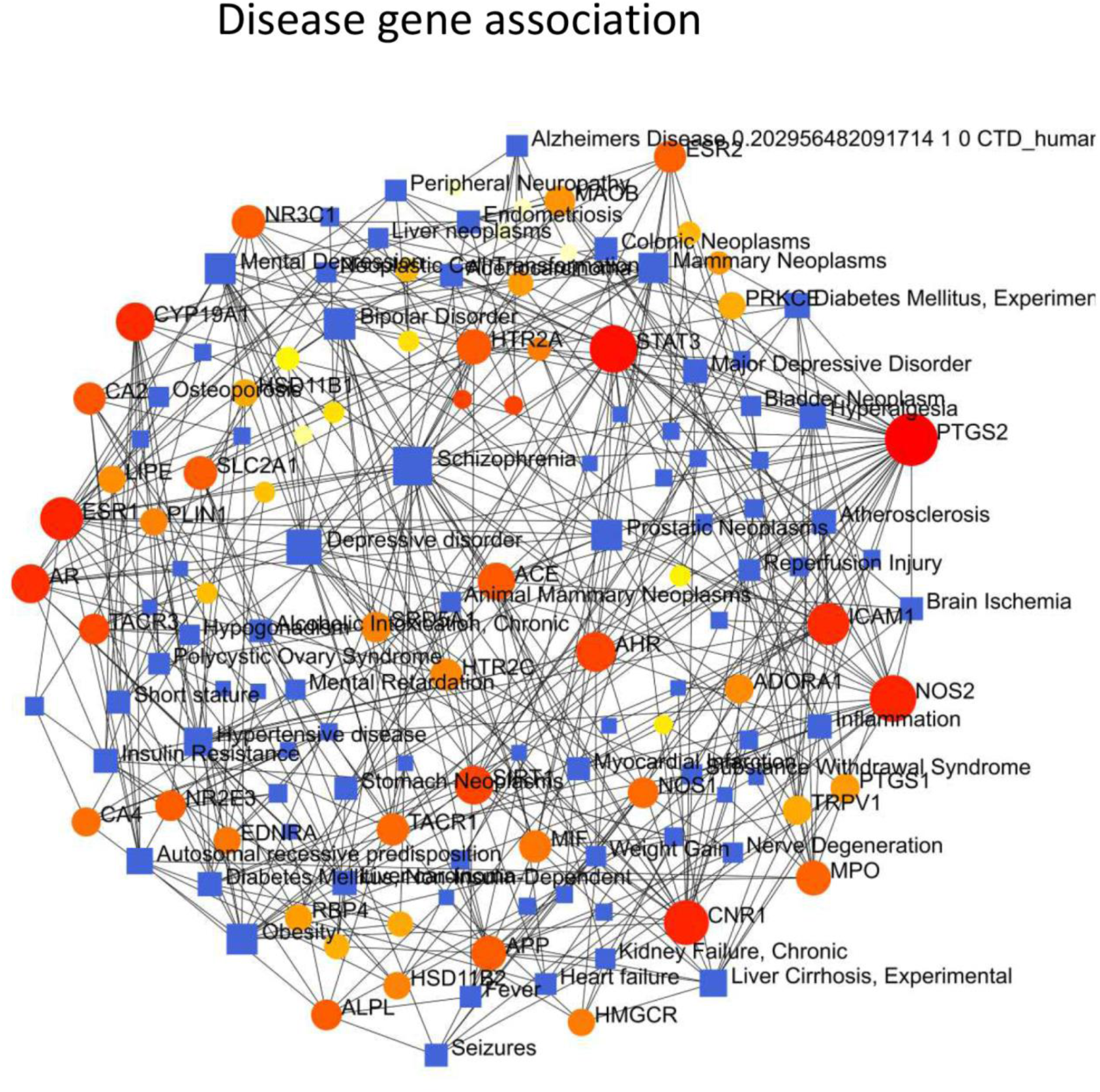
Gene-disease association network. Degree sorted network was constructed using Cytoscape v3.8.2. The nodes denote metabolites, targets, and pathways in the network, while the edges denote the interaction between the nodes. The node size and colour gradient (colour: red→ pink → purple; and size: large to small) represents the high to the low degree of the nodes within the network.

## Discussion

The present study investigated the elucidation of molecular mechanisms and therapeutic effects of *Trikatu*, an Indian polyherbal formulation utilizing an NP-based approach in T2DM, Obesity and Dyslipidemia. Screening of bioactive phytochemicals includes evaluation of DL,and ADME properties which mainly predict bioavailability, dynamic changes of drugs in the body and their half-life. Many compounds fail in drug development due to poor ADME properties (55). Our NP analysis showed that *Trikatu* contains twenty bioactive phytochemicals (E-beta-carotene, acetic acid, genistein, cholesterol formate, alpha-gurjunene, etc.), withpositive DL score and 30% OB. These bioactive phytochemicals were found to interact with several protein targets such as RXRA, ESR1, STAT3, GSK3B, TNF, APP, SIRT1, etc. which can modulate multiple biological pathways related to obesity, dyslipidemia, and T2DM.

### *Trikatu* bio-active phytochemicals regulate insulin resistance in T2DM and obesity

Both T2DM and obesity are chronic disorders with common pathophysiology (56–59). Insulin resistance (IR) is considered crucial in diabetes, obesity, and related complications (59–62). It refers to impaired insulin sensitivity to the biological response in the cell, subsequently affecting the glucose and lipid metabolism (60, 61). We found that genistein can regulate the insulin resistance pathway by targeting GSK3B, a kinase that regulates insulin-mediated glycogen metabolism by activating glycogen synthase (63). Genistein also protects pancreatic beta cells and improve insulin secretion, as well as regulate insulin resistance via the PI3K and AMPK pathways (63–66). Moreover, literature from other diseases like cancer provides empirical evidence that genistein regulates GSK3B in cell proliferation (67, 68). This study also supports the notion that genistein regulates GSK3B in the insulin resistance pathway. Similarly, piperlongumine has been shown to regulate STAT3 and TNF in the insulin resistance pathway. Our finding is consistent with previous studies which shows thatpiperlongumine modulates STAT3 and TNF in the regulation of insulin resistance in T2DM and obesity (69, 70).

### Trikatu bio-active phytochemicals regulate lipolysis, adipocytokine signalling in Dyslipidemia and Obesity

Patients with obesity and T2DM have high fasting and postprandial triglycerides, low HDL cholesterol, high LDL-cholesterol, inflammation, insulin resistance in fat cells, and fat accumulation in the liver (71). Additionally, the bile lipid and cholesterol secretion pathways are also affected in obesity and T2DM (72, 73). We found many bioactive phytochemicals of *Trikatu*, such as beta-carotene, acetic acid, beta-sitosterol, (3beta,5alpha)-Stigmastan-3-ol target proteins like RXRA, STAT-3, TNF, perilipin, and HSL which are critical for the regulation of lipid metabolism. Specifically, we found that E-beta caroteneregulates the adipocytokine signalling pathway by targeting RXRA, STAT-3, and TNF proteins. Previous studies support this, suggesting its preventive role in obesity and T2DM (74–76). Furthermore, another study has reported that asymmetric cleavage of E-beta carotene leads to the production of β-apo-14′-carotenal (apo14), which further interacts with RXRA to regulate adipogenesis (77).

Acetic acid targets the RXRA protein, further regulating the adipocytokine signalling pathway. Earlier studies have shown that acetic acid regulates lipid metabolism in the liver and skeletal muscles by regulating serum triglyceride, glucose, and insulin levels (78–80). In this study, we found that acetic acid regulates RXRA protein in the adipocyte signalling pathway, which is not previously documented, providing a clue to new targets of acetic acid in mediating lipid metabolism. Furthermore, we also found acetic acid mediating the lipolysis pathway through the involvement of hormonal lipase and perilipin-1. Hormonal lipases are enzymes that play an essential role in the hydrolysis of triglycerides, diacylglycerides, and cholesterol esters (81). Perilipin-1 regulates HSL (hormone-sensitive lipase) activity and thus plays a preventive role in lipolysis (82–84). NP analysis provides a valuable insight into the interaction of acetic acid with HSL and PLN1 in lipid metabolism, which has no direct evidence in previous literature.

We found that beta-sitosterol regulates de novo cholesterol-mediated steroidogenic apparatus, i.e., steroid hormone biosynthesis, thyroid hormone signalling pathway, and bile secretion by targeting proteins such as CYP19A1, ESR1, and NR1H4. Beta-sitosterol, a plant sterol with a similar structure to cholesterol, has been reported to have anti-diabetic, hypolipidemic, anti-cancer, anti-arthritic, and hepatoprotective properties (85, 86). Researches suggest that CYP19A1, one of the target proteins, converts C19 androgens to C18 estrogens (87). The ESR1 protein plays an essential role in estrogen-induced glucose homeostasis and thyroid metabolism (88, 89). Unlike previous literature, our study provides a novel clue to these interactions in steroid hormone synthesis. Stigmastanol is a reduction product of beta-sitosterol, which inhibits cholesterol absorption (90, 91). According to our results, Stigmastanol ((3beta,5alpha)-Stigmastan-3-ol) regulates NR1H4 in bile secretion. So this study provides a novel clue of these interactions in regulating bile secretion.

We further performed molecular docking to validate the interaction between target proteins and metabolites revealed in NP analysis. It provides a platform to explore ligand and receptor molecule conformations until the minimum binding energy is reached (92, 93). Based on the degree >10, proteins like ESR1, RXRA, and STAT3 with their respective ligands like piperlongumine, genistein, acetic acid, beta-sitosterol, cholesterol formate, ascorbic acid, 3-beta alpha stigmastanol, alpha gurjunene, beta-carotene etc. were selected for validation of our results. Our docking results were corroborated by the previous NP studies that showed stable binding of RXRA with E-beta carotene (94) and piperlongumine as a direct inhibitor of STAT3 in cancer (95). In addition, beta-sitosterol also exhibits stable binding to ESR1 in a study related to ulcerative colitis (96).

Our gene-disease association results revealed that phytochemicals of *Trikatu* target the proteins such as RXRA & STAT-3, which are implicated in diverse disease conditions, like depressive disorder, adenocarcinoma, hypertensive disease, atherosclerosis, inflammation, liver cirrhosis, fever, etc. In spite of their different causes, we observed that Trikatu, recommendations for treating them, reflected the holistic approach of Ayurvedic medicine based on the interconnectedness of metabolic pathways (15–17,19,21,97).

## Limitations

This study has several limitations. First, we have not experimentally validated the results through *in vivo* or *in vitro* studies. Second, due to limited data in databases, some of the protein targets of bioactive phytochemicals may be overlooked. Third, whether the pathways and protein targets obtained in our study were upregulated or downregulated is unclear. Fourth, some important phytochemicals like piperine and curcumin got excluded in the screening because they do not possess a positive DL score and 30% OB. Fifth, NP based analysis of traditional medicine does not have strict guidelines; however, a draft of NP was recently published, which summarizes software details, methods, and analysis should be mentioned while the presenting the study(98).

## Clinical implications

In the present study, we have used a reverse pharmacology and network pharmacology-based approach to identify the potential bioactive phytochemicals of *Trikatu* in treating metabolic diseases. It was found that *Trikatu* may play a crucial role in regulating insulin resistance and lipid metabolism via lipolysis and adipocytokine signaling, similar to conventional drugs against metabolic disorders. Therefore, this study may provide a basis for targeted preclinical and clinical studies investigating the efficacy and interactions of *Trikatu* in diabetic dyslipidemia and obesity. However, a deeper understanding of phytochemicals is needed to help customize dose and treatment protocol in clinical practice.

## Conclusion

The present study evaluated the bioactive phytochemicals of Trikatu, their targets, thereby elucidating its mechanism in treating T2DM, obesity, and dyslipidemia. Phytochemicals such as E-beta carotene, acetic acid, beta-sitosterol, genistein and piperlongumine act on proteins such as ESR1, STAT3, RXRA, GSK3B and TNF. These proteins regulate pathways related to glucose and lipid metabolism viz., insulin resistance, steroid hormone biosynthesis, regulation of lipolysis in adipocytes, adipocytokine & cGMP-PKG signalling pathways, arachidonic acid metabolism and bile secretion (Figure 8). The results suggest that Trikatu’s underlying mechanism against metabolic disorders is a synergy of multiple targets and multiple pathways. These findings may be further validated in future clinical studies against metabolic disorders.

**Figure 8.**
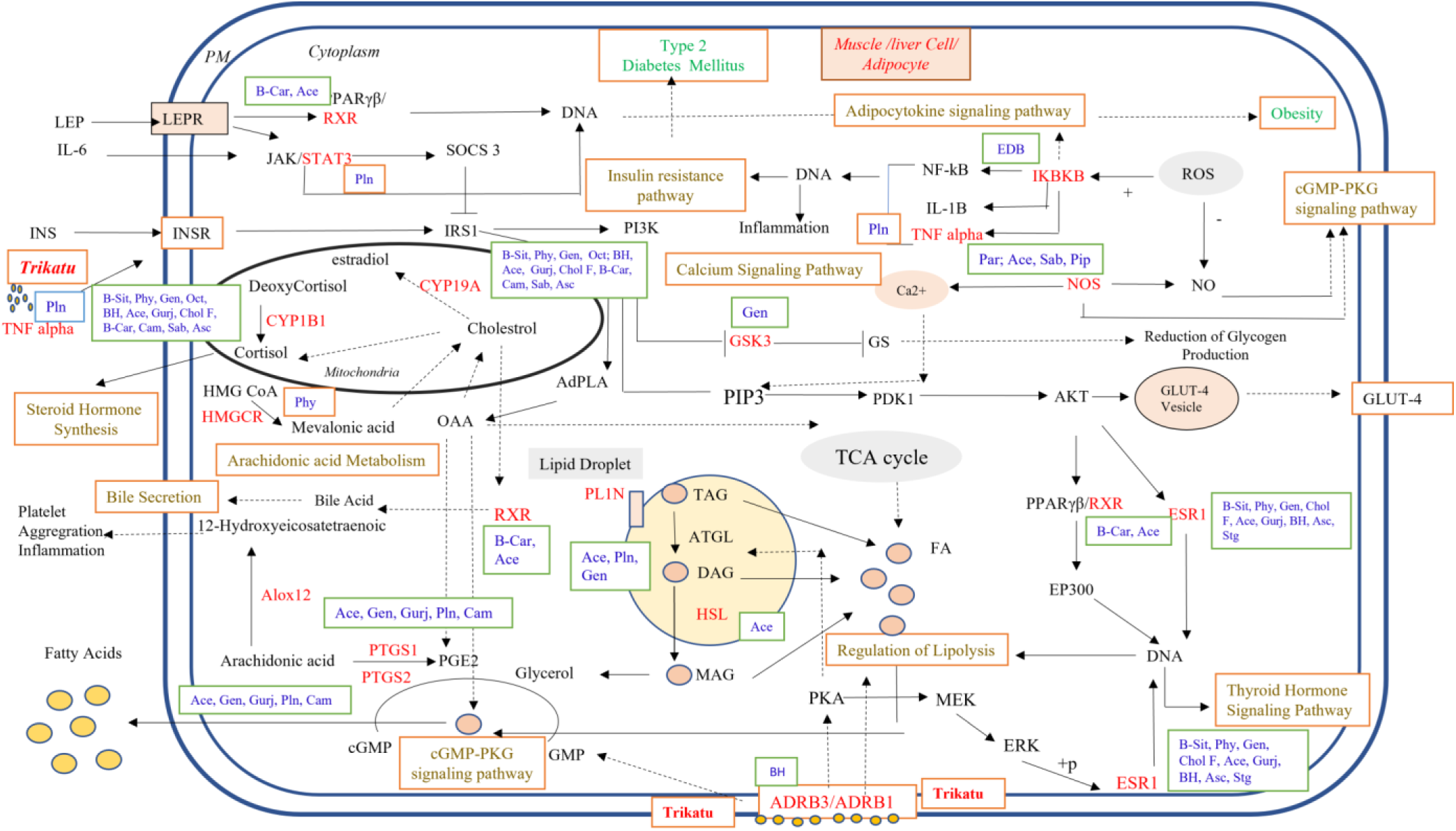
Schematic diagram showing targets and pathways regulated by *Trikatu* phytochemicals. *Tikatu* protein targets are shown in red and pathways are shown in brown. The phytochemicals are mainly shown in boxes: B-Sit: beta-sitosterol; Phy: phytosterol, Gen: genistein; Chol F: cholestrol formate; Ace: acetic acid; al-Gurj: alpha gurjunene, Asc: ascorbic acid; Stg: (3beta,5alpha)-Stigmastan-3-ol; Pln: piperlongumine; B-Car: beta carotene; Cam: camphor; EDB: 2-(Ethylamino)-4,5-dihydroxybenzamide; Gen: Genistein; Sab: (+)-Sabinene; PIP: Piperamide-C5:1(2E); Par: Paroxetine; Oct: 1-(4-Hydroxy-3-methoxyphenyl)-3,5-octanediol; BH: 1,7-Bis(4-hydroxy-3-methoxyphenyl)-3,5-heptanediol.

## Acknowledgements

The authors acknowledge the Ministry of AYUSH, Government of India, for their support through the Centre of Excellence (COE) scheme to the Centre for Integrative Medicine and Research (CIMR), AIIMS, New Delhi.

## Data Availability Statement

The data sets presented in this study can be found in online repositories. The names of the repository/repositories and accession number(s) can be found in the article/Supplementary Material.

## Supplementary Material

Supplementary data are available online.

## Author Contributions

VC : He has conceptualized, performed the experiments related to Network Pharmcology, interpreted and analysed the data, wrote original draft of the manuscript. MW : He has conceived and designed the study, analysed, performed the experiments related to Network Pharmcology, interpreted the data, review and edited the manuscript. AK : He has analysed, interpreted the data, and reviewed the manuscript. BKK : He has reviewed and edited the manuscript. VS : She has reviewed and edited the manuscript. SR : She has reviewed and edited the manuscript. PK : She has analyzed and interpreted the data. GS : He has supervised, conceived and designed the study, reviewed and edited the manuscript.

## Supplementary material

Supplementary data are available online

Supplementary File 1

Supplementary File 2

Supplementary File 3

Supplementary File 4

Supplementary File 5

## References

1. Stephens CR, Easton JF, Robles-Cabrera A, Fossion R, De la Cruz L, Martínez-Tapia R, Barajas-Martínez A, Hernández-Chávez A, López-Rivera JA, Rivera AL. The impact of education and age on metabolic disorders. Front public Heal (2020)180.

2. Wilson PWF, D’Agostino RB, Parise H, Sullivan L, Meigs JB. Metabolic syndrome as a precursor of cardiovascular disease and type 2 diabetes mellitus. Circulation (2005) 112:3066–3072.

3. Bonora E. The metabolic syndrome and cardiovascular disease. Ann Med (2006) 38:64–80.

4. Saklayen MG. The global epidemic of the metabolic syndrome. Curr Hypertens Rep (2018) 20:1–8.

5. Tabatabaei-Malazy O, Larijani B, Abdollahi M. Targeting metabolic disorders by natural products. J Diabetes Metab Disord (2015) 14:1–21.

6. Waltenberger B, Mocan A, Šmejkal K, Heiss EH, Atanasov AG. Natural products to counteract the epidemic of cardiovascular and metabolic disorders. Molecules (2016) 21: doi: 10.3390/molecules21060807

7. Chaudhury A, Duvoor C, Reddy Dendi VS, Kraleti S, Chada A, Ravilla R, Marco A, Shekhawat NS, Montales MT, Kuriakose K. Clinical review of antidiabetic drugs: implications for type 2 diabetes mellitus management. Front Endocrinol (Lausanne) (2017) 8:6.

8. Padhi S, Nayak AK, Behera A. Type II diabetes mellitus: A review on recent drug based therapeutics. Biomed Pharmacother (2020) 131:110708.

9. Sun Q, He M, Zhang M, Zeng S, Chen L, Zhao H, Yang H, Liu M, Ren S, Xu H. Traditional Chinese Medicine and Colorectal Cancer: Implications for Drug Discovery. Front Pharmacol (2021) 12:1557.

10. Nilashi M, Samad S, Yusuf SYM, Akbari E. Can complementary and alternative medicines be beneficial in the treatment of COVID-19 through improving immune system function? J Infect Public Health (2020) 13:893.

11. Debas HT, Laxminarayan R, Straus SE. Chapter 69: Complementary and Alternative Medicine. Dis Control Priorities Dev Ctries (2004)1281–1292.

12. Bodeker G, Ong C-K. WHO global atlas of traditional, complementary and alternative medicine. World Health Organization (2005).

13. Tipton CM, Tucson A, BUILDING INAG. HISTORICAL PERSPECTIVE: SUSRUTA OF INDIA, AN UNRECOGNIZED CONTRIBUTOR TO THE HISTORY OF EXERCISE PHYSIOLOGY. (2008)

14. Iyer A, Panchal S, Poudyal H, Brown L. Potential health benefits of Indian spices in the symptoms of the metabolic syndrome: a review. (2009)

15. Kaushik R, Jain J, Khan AD. “Trikatu: Transforming Food into Medicines.,” Innovations in Food Technology. Springer (2020). p. 501–508

16. Kaushik R. Trikatu-A combination of three bioavailability enhancers. Int J Green Pharm (2018) 12:

17. Govindarajan N, Chinnapillai A, Balasundaram M, Narasimhaji CV, Ganji K, Raju I. Pharmacognostical and Phytochemical Evaluation of a Polyherbal Ayurvedic Formulation Trikatu Churna. J Ayurveda Med Sci (2016) 1:

18. Sivakumar V, Sivakumar S. Effect of an indigenous herbal compound preparation ‘Trikatu’on the lipid profiles of atherogenic diet and standard diet fed Rattus norvegicus. Phyther Res (2004) 18:976–981.

19. Johri RK, Zutshi U. An Ayurvedic formulation ‘Trikatu’and its constituents. J Ethnopharmacol (1992) 37:85–91.

20. Kumawat VB, Sharma SK, Sharma UK, Sharma S. Effect of trikatu compound in hypercholesteremia-a clinical study. Environ Conserv J (2015) 16:57–61.

21. Krishniya K, Sharma K, Padhar BKC, Joshi RK. A therapeutic review on Trikatu Churna in the management of Sthaulya (Obesity). J Ayurveda Integr Med Sci (2020) 5:258–261.

22. Hopkins AL. Network pharmacology: the next paradigm in drug discovery. Nat Chem Biol (2008) 4:682–690.

23. Wadhawan M, Chhabra V, Katiyar A, Sharma V, Khuntia BK, Rathore S, Kaur P, Sharma G. A Network Pharmacology-Based Approach to Explore Therapeutic Mechanism of Indian Herbal Formulation Nisha Amalaki in Treating Type 2 Diabetes Mellitus. (2021)

24. Hopkins AL. Network pharmacology. Nat Biotechnol (2007) 25:1110–1111.

25. Chandran U, Mehendale N, Patil S, Chaguturu R, Patwardhan B. Network pharmacology. Innov Approaches Drug Discov (2017)127.

26. Kim S, Chen J, Cheng T, Gindulyte A, He J, He S, Li Q, Shoemaker BA, Thiessen PA, Yu B. PubChem 2019 update: improved access to chemical data. Nucleic Acids Res (2019) 47:D1102–D1109.

27. Afendi FM, Okada T, Yamazaki M, Hirai-Morita A, Nakamura Y, Nakamura K, Ikeda S, Takahashi H, Altaf-Ul-Amin M, Darusman LK. KNApSAcK family databases: integrated metabolite–plant species databases for multifaceted plant research. Plant Cell Physiol (2012) 53:e1–e1.

28. Mohanraj K, Karthikeyan BS, Vivek-Ananth RP, Chand RPB, Aparna SR, Mangalapandi P, Samal A. IMPPAT: A curated database of Indian medicinal plants, phytochemistry and therapeutics. bioRxiv (2017)206995.

29. Degtyarenko K, De Matos P, Ennis M, Hastings J, Zbinden M, McNaught A, Alcántara R, Darsow M, Guedj M, Ashburner M. ChEBI: a database and ontology for chemical entities of biological interest. Nucleic Acids Res (2007) 36:D344–D350.

30. Wang Z, Liang L, Yin Z, Lin J. Improving chemical similarity ensemble approach in target prediction. J Cheminform (2016) 8:1–10.

31. Dong J, Wang N-N, Yao Z-J, Zhang L, Cheng Y, Ouyang D, Lu A-P, Cao D-S. ADMETlab: a platform for systematic ADMET evaluation based on a comprehensively collected ADMET database. J Cheminform (2018) 10:1–11.

32. Khanal P, Duyu T, Patil BM, Dey YN, Pasha I, Wanjari M, Gurav SS, Maity A. Network pharmacology of AYUSH recommended immune-boosting medicinal plants against COVID-19. J Ayurveda Integr Med (2020)

33. Cheng F, Li W, Zhou Y, Shen J, Wu Z, Liu G, Lee PW, Tang Y. admetSAR: a comprehensive source and free tool for assessment of chemical ADMET properties. (2012)

34. Keiser MJ, Roth BL, Armbruster BN, Ernsberger P, Irwin JJ, Shoichet BK. Relating protein pharmacology by ligand chemistry. Nat Biotechnol (2007) 25:197–206. doi: 10.1038/nbt1284

35. Yao Z-J, Dong J, Che Y-J, Zhu M-F, Wen M, Wang N-N, Wang S, Lu A-P, Cao D-S. TargetNet: a web service for predicting potential drug–target interaction profiling via multi-target SAR models. J Comput Aided Mol Des (2016) 30:413–424.

36. Gfeller D, Grosdidier A, Wirth M, Daina A, Michielin O, Zoete V. SwissTargetPrediction: a web server for target prediction of bioactive small molecules. Nucleic Acids Res (2014) 42:W32–W38.

37. Piñero J, Bravo À, Queralt-Rosinach N, Gutiérrez-Sacristán A, Deu-Pons J, Centeno E, García-García J, Sanz F, Furlong LI. DisGeNET: a comprehensive platform integrating information on human disease-associated genes and variants. Nucleic Acids Res (2016)gkw943.

38. Hamosh A, Scott AF, Amberger JS, Bocchini CA, McKusick VA. Online Mendelian Inheritance in Man (OMIM), a knowledgebase of human genes and genetic disorders. Nucleic Acids Res (2005) 33:D514–D517.

39. Wang Y, Zhang S, Li F, Zhou Y, Zhang Y, Wang Z, Zhang R, Zhu J, Ren Y, Tan Y. Therapeutic target database 2020: enriched resource for facilitating research and early development of targeted therapeutics. Nucleic Acids Res (2020) 48:D1031–D1041.

40. Kanehisa M, Goto S, Furumichi M, Tanabe M, Hirakawa M. KEGG for representation and analysis of molecular networks involving diseases and drugs. Nucleic Acids Res (2010) 38:D355–D360.

41. Consortium U. UniProt: a hub for protein information. Nucleic Acids Res (2015) 43:D204–D212.

42. Oliveros JC. Venny 2.1. 0. Venny An Interact Tool Comp List with Venn’s Diagrams(2007-2015) Available online http//bioinfogp cnb csic es/tools/venny/(Accessed Febr 15, 2016) (2016)

43. Zhou G, Soufan O, Ewald J, Hancock REW, Basu N, Xia J. NetworkAnalyst 3.0: a visual analytics platform for comprehensive gene expression profiling and meta-analysis. Nucleic Acids Res (2019) 47:W234–W241.

44. Breuer K, Foroushani AK, Laird MR, Chen C, Sribnaia A, Lo R, Winsor GL, Hancock REW, Brinkman FSL, Lynn DJ. InnateDB: Systems biology of innate immunity and beyond - Recent updates and continuing curation. Nucleic Acids Res (2013) 41:1228–1233. doi: 10.1093/nar/gks1147

45. Shannon P, Markiel A, Ozier O, Baliga NS, Wang JT, Ramage D, Amin N, Schwikowski B, Ideker T. Cytoscape: a software environment for integrated models of biomolecular interaction networks. Genome Res (2003) 13:2498–2504.

46. Dennis G, Sherman BT, Hosack DA, Yang J, Gao W, Lane HC, Lempicki RA. DAVID: database for annotation, visualization, and integrated discovery. Genome Biol (2003) 4:1–11.

47. Wickham H. ggplot2. Wiley Interdiscip Rev Comput Stat (2011) 3:180–185.

48. Wickham H, Chang W, Henry L. ggplot2. Comput software] Retrieved from http//ggplot2 org (2012)

49. Skern T. “An Archive and a Tool: PDB and PyMOL.,” Exploring Protein Structure: Principles and Practice. Springer (2018). p. 7–28

50. Tian W, Chen C, Lei X, Zhao J, Liang J. CASTp 3.0: computed atlas of surface topography of proteins. Nucleic Acids Res (2018) 46:W363–W367.

51. Huey R, Morris GM, Forli S. Using AutoDock 4 and AutoDock vina with AutoDockTools: a tutorial. Scripps Res Inst Mol Graph Lab (2012) 10550:92037.

52. Trott O, Olson AJ. AutoDock Vina: improving the speed and accuracy of docking with a new scoring function, efficient optimization, and multithreading. J Comput Chem (2010) 31:455–461.

53. Salentin S, Schreiber S, Haupt VJ, Adasme MF, Schroeder M. PLIP: fully automated protein–ligand interaction profiler. Nucleic Acids Res (2015) 43:W443–W447.

54. Laskowski RA, Swindells MB. LigPlot+: multiple ligand–protein interaction diagrams for drug discovery. (2011)

55. Zhang W, Huai Y, Miao Z, Qian A, Wang Y. Systems pharmacology for investigation of the mechanisms of action of traditional Chinese medicine in drug discovery. Front Pharmacol (2019) 10:743.

56. Yoon K-H, Lee J-H, Kim J-W, Cho JH, Choi Y-H, Ko S-H, Zimmet P, Son H-Y. Epidemic obesity and type 2 diabetes in Asia. Lancet (2006) 368:1681–1688.

57. Sadry SA, Drucker DJ. Emerging combinatorial hormone therapies for the treatment of obesity and T2DM. Nat Rev Endocrinol (2013) 9:425–433.

58. Rochlani Y, Pothineni NV, Kovelamudi S, Mehta JL. Metabolic syndrome: pathophysiology, management, and modulation by natural compounds. Ther Adv Cardiovasc Dis (2017) 11:215–225.

59. Maggio CA, Pi-Sunyer FX. The prevention and treatment of obesity: application to type 2 diabetes. Diabetes Care (1997) 20:1744–1766.

60. Petersen MC, Shulman GI. Mechanisms of insulin action and insulin resistance. Physiol Rev (2018) 98:2133–2223.

61. Wilcox G. Insulin and insulin resistance. Clin Biochem Rev (2005) 26:19.

62. Malone JI, Hansen BC. Does obesity cause type 2 diabetes mellitus (T2DM)? Or is it the opposite? Pediatr Diabetes (2019) 20:5–9.

63. Arunkumar E, Karthik D, Anuradha CV. Genistein sensitizes hepatic insulin signaling and modulates lipid regulatory genes through p70 ribosomal S6 kinase-1 inhibition in high-fatâ€“high-fructose diet-fed mice. Pharm Biol (2013) 51:815–824.

64. Li R, Ding X-W, Geetha T, Al-Nakkash L, Broderick TL, Babu JR. Beneficial Effect of Genistein on Diabetes-Induced Brain Damage in the ob/ob Mouse Model. Drug Des Devel Ther (2020) 14:3325.

65. Arunkumar E, Anuradha CV. Genistein promotes insulin action through adenosine monophosphate–activated protein kinase activation and p70 ribosomal protein S6 kinase 1 inhibition in the skeletal muscle of mice fed a high energy diet. Nutr Res (2012) 32:617–625.

66. Behloul N, Wu G. Genistein: a promising therapeutic agent for obesity and diabetes treatment. Eur J Pharmacol (2013) 698:31–38.

67. El Touny LH, Banerjee PP. Akt–GSK-3 pathway as a target in genistein-induced inhibition of TRAMP prostate cancer progression toward a poorly differentiated phenotype. Carcinogenesis (2007) 28:1710–1717.

68. Erten F, Yenice E, Orhan C, Er B, Öner PD, Deeh PBD, Şahin K. Genistein suppresses the inflammation and GSK-3 pathway in an animal model of spontaneous ovarian cancer. Turkish J Med Sci (2021) 51:1465–1471.

69. Xu P, Xiao J, Chi S. Piperlongumine attenuates oxidative stress, inflammatory, and apoptosis through modulating the GLUT-2/4 and AKT signaling pathway in streptozotocin-induced diabetic rats. J Biochem Mol Toxicol (2021) 35:1–12.

70. Chen Q, Lv J, Yang W, Xu B, Wang Z, Yu Z, Wu J, Yang Y, Han Y. Targeted inhibition of STAT3 as a potential treatment strategy for atherosclerosis. Theranostics (2019) 9:6424.

71. Wu L, Parhofer KG. Diabetic dyslipidemia. Metabolism (2014) 63:1469–1479.

72. Li T, Chiang JYL. Bile acid signaling in metabolic disease and drug therapy. Pharmacol Rev (2014) 66:948–983.

73. Ahmad TR, Haeusler RA. Bile acids in glucose metabolism and insulin signalling— mechanisms and research needs. Nat Rev Endocrinol (2019) 15:701–712.

74. Kameji H, Mochizuki K, Miyoshi N, Goda T. β-Carotene accumulation in 3T3-L1 adipocytes inhibits the elevation of reactive oxygen species and the suppression of genes related to insulin sensitivity induced by tumor necrosis factor-α. Nutrition (2010) 26:1151–1156.

75. Cho SO, Kim M-H, Kim H. β-Carotene inhibits activation of NF-κB, activator protein-1, and STAT3 and regulates abnormal expression of some adipokines in 3T3-L1 adipocytes. J cancer Prev (2018) 23:37.

76. Marcelino G, Machate DJ, Freitas K de C, Hiane PA, Maldonade IR, Pott A, Asato MA, Candido CJ, GuimarÃ£es R de CA. Beta-Carotene: Preventive Role for Type 2 Diabetes Mellitus and Obesity: A Review. Molecules (2020) 25:5803.

77. Ziouzenkova O, Orasanu G, Sukhova G, Lau E, Berger JP, Tang G, Krinsky NI, Dolnikowski GG, Plutzky J. Asymmetric cleavage of β-carotene yields a transcriptional repressor of retinoid X receptor and peroxisome proliferator-activated receptor responses. Mol Endocrinol (2007) 21:77–88.

78. Yamashita H. Biological function of acetic acid–improvement in obesity and glucose tolerance by acetic acid in type 2 diabetic rats. Crit Rev Food Sci Nutr (2016) 56:S171–S175.

79. Mitrou P, Petsiou E, Papakonstantinou E, Maratou E, Lambadiari V, Dimitriadis P, Spanoudi F, Raptis SA, Dimitriadis G. The role of acetic acid on glucose uptake and blood flow rates in the skeletal muscle in humans with impaired glucose tolerance. Eur J Clin Nutr (2015) 69:734–739.

80. Santos HO, de Moraes WMAM, da Silva GAR, Prestes J, Schoenfeld BJ. Vinegar (acetic acid) intake on glucose metabolism: A narrative review. Clin Nutr ESPEN (2019) 32:1–7.

81. Kraemer FB, Shen W-J. Hormone-sensitive lipase. J Lipid Res (2002) 43:1585–1594.

82. Kim S-J, Tang T, Abbott M, Viscarra JA, Wang Y, Sul HS. AMPK phosphorylates desnutrin/ATGL and hormone-sensitive lipase to regulate lipolysis and fatty acid oxidation within adipose tissue. Mol Cell Biol (2016) 36:1961–1976.

83. Sztalryd C, Xu G, Dorward H, Tansey JT, Contreras JA, Kimmel AR, Londos C. Perilipin A is essential for the translocation of hormone-sensitive lipase during lipolytic activation. J Cell Biol (2003) 161:1093–1103.

84. Blanchette-Mackie EJ, Dwyer NK, Barber T, Coxey RA, Takeda T, Rondinone CM, Theodorakis JL, Greenberg AS, Londos C. Perilipin is located on the surface layer of intracellular lipid droplets in adipocytes. J Lipid Res (1995) 36:1211–1226.

85. Jayaraman S, Devarajan N, Rajagopal P, Babu S, Ganesan SK, Veeraraghavan VP, Palanisamy CP, Cui B, Periyasamy V, Chandrasekar K. β-Sitosterol Circumvents Obesity Induced Inflammation and Insulin Resistance by down-Regulating IKKβ/NF- κB and JNK Signaling Pathway in Adipocytes of Type 2 Diabetic Rats. Molecules (2021) 26:2101.

86. Gumede NM, Lembede BW, Brooksbank RL, Erlwanger KH, Chivandi E. β-Sitosterol shows potential to protect against the development of high-fructose diet-induced metabolic dysfunction in female rats. J Med Food (2020) 23:367–374.

87. Boon WC, Chow JDY, Simpson ER. The multiple roles of estrogens and the enzyme aromatase. Prog Brain Res (2010) 181:209–232.

88. Fatima LA, Campello RS, de Souza Santos R, Freitas HS, Frank AP, Machado UF, Clegg DJ. Estrogen receptor 1 (ESR1) regulates VEGFA in adipose tissue. Sci Rep (2017) 7:1–14.

89. Handgraaf S, Riant E, Fabre A, Waget A, Burcelin R, Lière P, Krust A, Chambon P, Arnal J-F, Gourdy P. Prevention of obesity and insulin resistance by estrogens requires ERα activation function-2 (ERαAF-2), whereas ERαAF-1 is dispensable. Diabetes (2013) 62:4098–4108.

90. Heinemann T, Kullak-Ublick G-A, Pietruck B, Von Bergmann K. Mechanisms of action of plant sterols on inhibition of cholesterol absorption. Eur J Clin Pharmacol (1991) 40:S59–S63.

91. Batta AK, Xu G, Honda A, Miyazaki T, Salen G. Stigmasterol reduces plasma cholesterol levels and inhibits hepatic synthesis and intestinal absorption in the rat. Metabolism (2006) 55:292–299.

92. Pagadala NS, Syed K, Tuszynski J. Software for molecular docking: a review. Biophys Rev (2017) 9:91–102.

93. Ferreira LG, Dos Santos RN, Oliva G, Andricopulo AD. Molecular docking and structure-based drug design strategies. Molecules (2015) 20:13384–13421.

94. Huang M-Y, Wang Z-Z, Long J-L, Yang X-Y, Zhang Y, Yan D. Mechanism of Jinqi Jiangtang Tablets in treatment of pancreatic β cell dysfunction based on network pharmacology and molecular docking technology. Zhongguo Zhong yao za zhi= Zhongguo Zhongyao Zazhi= China J Chinese Mater Medica (2021) 46:5341–5350.

95. Bharadwaj U, Eckols TK, Kolosov M, Kasembeli MM, Adam A, Torres D, Zhang X, Dobrolecki LE, Wei W, Lewis MT. Drug-repositioning screening identified piperlongumine as a direct STAT3 inhibitor with potent activity against breast cancer. Oncogene (2015) 34:1341–1353.

96. Wang B, Liu Y, Sun J, Zhang N, Zheng X, Liu Q. Exploring the Potential Mechanism of Xiaokui Jiedu Decoction for Ulcerative Colitis Based on Network Pharmacology and Molecular Docking. J Healthc Eng (2021) 2021:

97. Rath C, Mane SS, Mangal A, Joseph GVR, Kumar A. “A Review on Trikatu an Ayurvedic formulation with special reference to.

98. Li S. Network pharmacology evaluation method guidance-draft. World J Tradit Chinese Med (2021) 7:146.

